# Connecting sequence features within the disordered C-terminal linker of *B. subtilis* FtsZ to functions and bacterial cell division

**DOI:** 10.1101/2022.06.29.498098

**Authors:** Min Kyung Shinn, Megan C. Cohan, Jessie L. Bullock, Kiersten M. Ruff, Petra A. Levin, Rohit V. Pappu

## Abstract

Intrinsically disordered regions (IDRs) can function as autoregulators of folded enzymes to which they are tethered. One example is the bacterial cell division protein, FtsZ. This includes a folded core and a C-terminal tail (CTT) that encompasses a poorly conserved, disordered C-terminal linker (CTL) and a well-conserved 17-residue C-terminal peptide (CT17). Sites for GTPase activity of FtsZs are formed at the interface between GTP binding sites and T7 loops on cores of adjacent subunits within dimers. Here, we explore the basis of autoregulatory functions of the CTT in *Bacillus subtilis* FtsZ (*Bs-*FtsZ). Molecular simulations show that the CT17 of *Bs-* FtsZ makes statistically significant CTL-mediated contacts with the T7 loop. Statistical Coupling Analysis of more than 10^3^ sequences from FtsZ orthologs reveals clear covariation of the T7 loop and the CT17 with most of the core domain whereas the CTL is under independent selection. Despite this, we discover the conservation of non-random sequence patterns within CTLs across orthologs. To test how the non-random patterns of CTLs mediate CTT-core interactions and modulate FtsZ functionalities, we designed *Bs-*FtsZ variants by altering the patterning of oppositely charged residues within the CTL. Such alterations disrupt the core-CTT interactions, lead to anomalous assembly and inefficient GTP hydrolysis *in vitro* and protein degradation, aberrant assembly, and disruption of cell division *in vivo*. Our findings suggest that viable CTLs in FtsZs are likely to be IDRs that encompass non-random, functionally relevant sequence patterns that also preserve three-way covariation of the CT17, the T7 loop, and core domain.

**Significance Statement:** Z-ring formation by the protein FtsZ controls cell division in rod-shaped bacteria. The C-terminus of FtsZ encompasses a disordered C-terminal linker (CTL) and a conserved CT17 motif. Both modules are essential for Z-ring formation and proper localization of FtsZ in cells. Previous studies suggested that generic intrinsically disordered regions (IDRs) might be suitable functional replacements for naturally occurring CTLs. Contrary to this suggestion, we find that the sequence-encoded conformational properties of CTLs help mediate autoregulatory interactions between covarying regions within FtsZ. Functional properties of the CTL are encoded via evolutionarily conserved, non-random sequence patterns. Disruption of these patterns impair molecular functions and cellular phenotypes. Our findings have broad implications for discovering functionally consequential sequence features within IDRs of other proteins.

## Introduction

Intrinsically disordered regions contribute to a multitude of protein functions (1-4). A common occurrence is of autoregulatory IDRs tethered either as tails to folded domains or as linkers between folded domains (5-12). Of particular interest are IDRs tethered to folded domains that are enzymes (7, 13, 14). Several studies demonstrate that IDRs tethered to folded domains can function as autoregulators (12), specifically as autoinhibitors of enzymatic activities (13, 15, 16). One such example is the C-terminal tail (CTT) of the essential GTPase that controls and regulates bacterial cell division (17). The CTT encompasses a disordered C-terminal linker (CTL) and an alpha-helix-forming C-terminal peptide.

Cell division in bacteria is initiated by assembly of the cytokinetic ring at the nascent division site (18-26). Polymers formed by the essential GTPase **F**ilamenting **t**emperature-**s**ensitive mutant **Z** (FtsZ) are the foundation of this ring, which is also known as the Z-ring (27-32). FtsZ is a prokaryotic homolog of tubulin. It forms single-stranded protofilaments upon binding GTP *in vitro* (33). Linear polymers of FtsZ, which also undergo bundling via lateral associations, serve as a platform for the cell division machinery composed of at least thirty different proteins (19, 32, 34-39). FtsZ polymers also undergo treadmilling *in vivo*, driven by the turnover of subunits that occurs on the order of seconds (40).

Previous *in vitro* experiments showed that FtsZ polymerization belongs to a class of phase transitions known as reversible polymerization (41). A defining hallmark of reversible phase transitions, with subunit concentration as the conserved order parameter, is the presence of at least one threshold concentration for the occurrence of a specific phase transition. Cohan et al., recently identified two distinct threshold concentrations for *B. subtilis* FtsZ (*Bs*-FtsZ) phase transitions occurring in the presence of GTP (17). In agreement with previous work on *E. coli* FtsZ, *Bs*-FtsZ forms single-stranded protofilaments when the first threshold concentration, denoted as *c*_A_, is crossed (42-47). The second threshold concentration, denoted as *c*_B_ where *c*_B_ > *c*_A_, characterizes the threshold for bundling of protofilaments.

*Bs-*FtsZ encompasses two domains: a folded N-terminal core and a C-terminal tail (CTT) (**Figure 1A**). The core domain forms a complete GTPase upon dimerization whereby the T7 loop of one protomer is inserted into the nucleotide binding site of the complementary protomer. The interface between the T7 loop and the nucleotide binding site is the active site for GTP hydrolysis (48). The CTT is further composed of an intrinsically disordered C-terminal linker (CTL) and a 17-residue C-terminal peptide (CT17). The CT17 was previously termed CTP (17). It includes a conserved “constant region” and “variable region (CTV)” (30, 49, 50). CT17 can form alpha-helical conformations (51-53) and is thus an alpha-molecular recognition element (54) that enables a precise network of homotypic and heterotypic protein-protein interactions. Whereas CT17 includes a conserved region (33), the CTL is hypervariable across orthologs, varying in length, amino acid composition, and sequence (33, 49, 55, 56). Mutations in the CTL and the CTV of *Bs*-FtsZ disrupt lateral interactions between protofilaments (49, 56).

**Figure 1:**
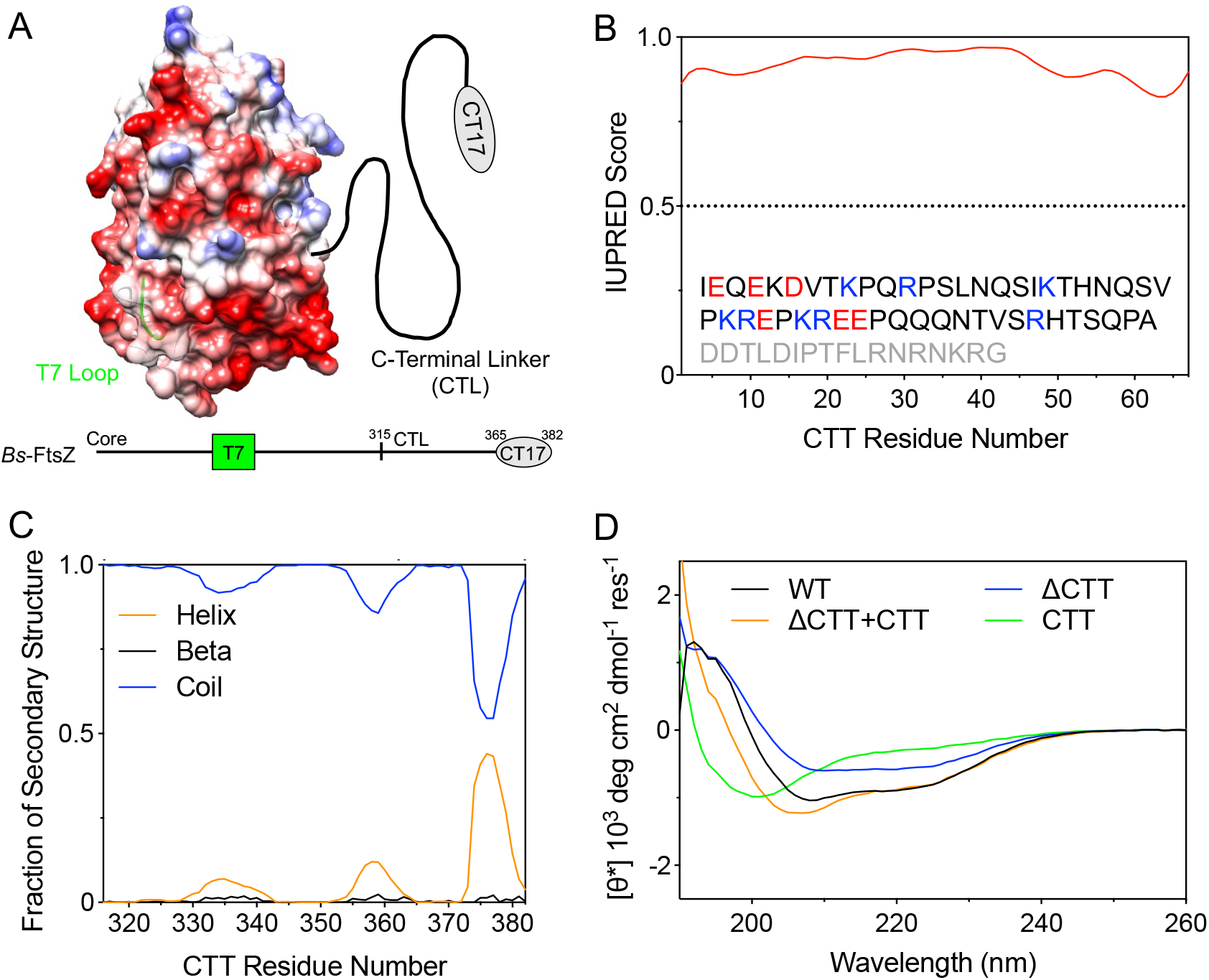
Modular architecture of *B. subtilis* FtsZ includes a disordered CTT. (A) The electrostatic potential (63) is mapped onto the core domain in red and blue for regions of negative and positive potential, respectively. The T7 loop is highlighted in green. The CTT includes a C-terminal linker (CTL) that connects the 17-residue C-terminal peptide (CT17) to the core domain. (B) The CTT is predicted to be disordered using IUPRED (60). The CTT sequence is shown with negatively and positively charged residues of the CTL in red and blue, respectively. The CT17 sequence is shown in gray. (C) Ensemble-averaged secondary structure contents of the CTT obtained from atomistic simulations. (D) UV-CD spectra of the four *Bs*-FtsZ constructs. See Methods for details on scaling of the [θ*].

Consistent with previous work (57-59), Cohan et al., showed, through systematic deletions of each module, that the core domain of *Bs*-FtsZ is the main driver of GTP binding induced polymerization (17). Deletion of the CT17 (ΔCT17), previously referred to as ΔCTP, increases *c*_A_ while also shifting *c*_B_ upward by at least three-fold. Internal deletion of the CTL (ΔCTL) decreases *c*_A_ and this construct forms mini rings stabilized by cohesive interactions of the CT17. Overall, the CTL weakens the driving forces for linear polymerization and bundling, whereas the CT17 appears to be the primary driver of lateral associations. Deletion of the CTT (ΔCTT) lowers *c*_A_ by over an order of magnitude and forms long, single-stranded polymers. Cohan et al., also showed that ΔCTT is the most efficient GTPase, whereas the wild-type *Bs*-FtsZ is the least efficient enzyme of the four constructs studied (17).

The picture that emerges is of the CTT as an autoregulator of *Bs*-FtsZ assembly and an autoinhibitor of enzymatic activity (17). Here, we uncover a molecular-level, mechanistic understanding of how the distinctive functions of CTTs are achieved.

## Results

### The CTL and CT17 interact with the core domain at mutually exclusive sites

The CTT is predicted to be disordered with IUPRED scores being above 0.5 for the entirety of the CTT (**Figure 1B**) (30, 60). We performed atomistic simulations to map conformational preferences of the isolated CTT peptide from *Bs-*FtsZ. These simulations show that the CTT of *Bs-*FtsZ samples a heterogenous ensemble of conformations with alpha-helical and random-coil character (**Figure 1C**). A large fraction of the alpha-helical signal arises from the CT17 suggesting that it is an example of an alpha-helical molecular recognition element / feature (α-MoRE or α-MoRF) (61, 62). Consistent with the predictions from simulations, ultraviolet circular dichroism (UV-CD) measurements for the isolated CTT peptide indicate that it is conformationally heterogeneous (Figs. **1D**, *SI Appendix*, **Figure S1**).

The CD spectrum for the ΔCTT construct, which refers to the core domain without the CTT, is consistent with a mixture of structures with minima at 208 and 222 nm, indicating the presence of alpha-helical and beta-sheet structures. The spectrum of full-length FtsZ closely resembles that of the ΔCTT construct with a mixture of structures. The sum of the spectra for the core domain and the CTT does not reproduce the spectrum of a full-length protein. This suggests that the CTT likely undergoes conformational changes when it is tethered to the core domain by forming inter-module interactions with the core.

We also performed atomistic simulations of the full-length wild-type (WT) *Bs-*FtsZ protomer to investigate the presence and types of interactions among the core domain, the CTL, and the CT17. The core-CTL and core-CT17 interactions were defined by the presence of residues within the CTL or CT17 being within 10 Å of a core residue. The frequencies of contacts were calculated from simulations. These are shown as a heatmap that maps the contacts being made onto the structure of the core domain. The CT17 fluctuates into and out of making contacts with the T7 loop (**Figure 2A**), whereas contacts between the CTL and the core exclude the T7 loop and mainly involve residues that are spatially proximal to the region on the core where the CTT is N-terminally tethered (**Figure 2C**). Synergies between the core-CTL and core-CT17 interactions were definitively unmasked in simulations of the different deletion constructs *viz*., the ΔCTL and ΔCT17 that were studied by Cohan et al., (17). The CT17 does not interact with the T7 loop when the CTL is deleted. Instead, alternate contacts form with the residues that are spatially proximal to the site where the CT17 is N-terminally tethered to the core domain in the ΔCTL construct (**Figure 2B**). Conversely, when the CT17 is deleted, the interactions between the CTL and the core that form in the full-length WT *Bs*-FtsZ protomer are preserved. Additionally, there are statistically significant contacts between the CTL and the T7 loop in the absence of CT17.

**Figure 2:**
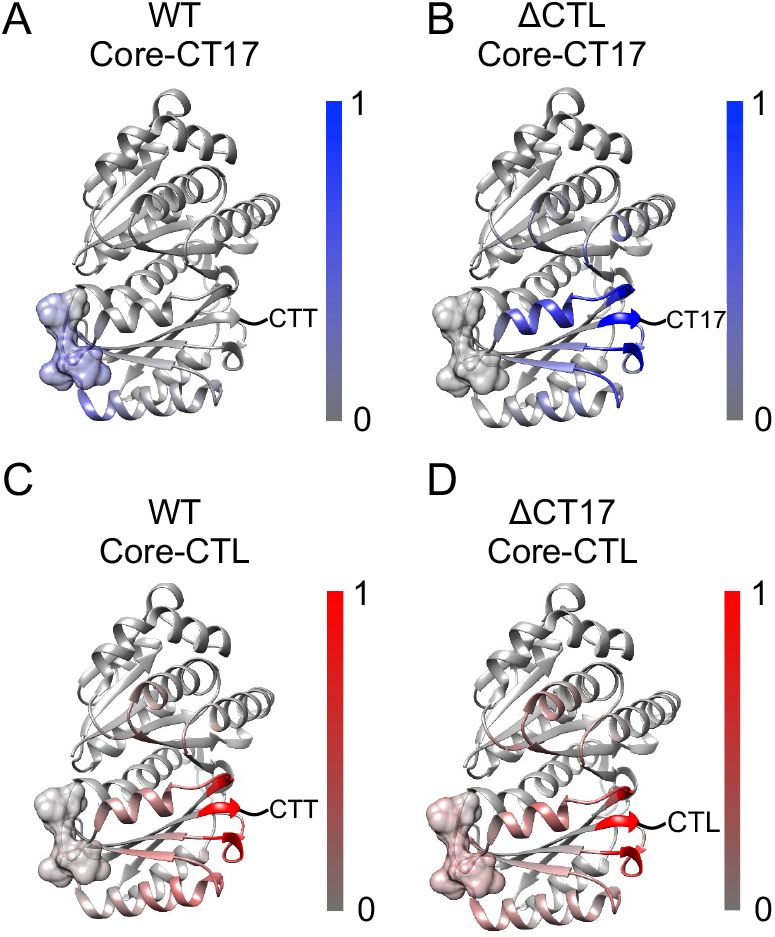
The CT17 and the CTL interact with mutually exclusive sites on the surface of the core domain. Color bars show the contact frequency between the core and (A, B) CT17 (blue) or (C, D) CTL (red) in WT, ΔCTL, and ΔCT17 constructs. The T7 loop is shown in space-filling model. The C-terminus of the core domain where the CTT emanates is noted.

Overall, the simulations show that the CTL and the CT17 can compete for interactions with the T7 loop of the core domain. However, in the full-length protomer, interactions between the CT17 and the T7 loop are favored over those with the CTL. Thus, the CTL becomes the facilitator of distal contacts between the CT17 and the T7 loop of the core domain. These results suggest the presence of “on” and “off” states whereby in the “on” state, the CT17 occupies the T7 loop, and in the “off” state, the CT17 is dissociated from the T7 loop. Importantly, as discussed in the *SI Appendix* (*SI Appendix*, **Figure S2**), the CTL-mediated contacts cannot be explained purely based on the intrinsic flexibility of the disordered CTL. Instead, these contacts appear to be sequence specific. Accordingly, we investigated the evolutionary basis for sequence-specific, CTL-mediated contacts.

### The CTL is under independent selection whereas the T7 loop and CTP belong to the same covarying sector

While the core domain and the CT17 are conserved, the CTL is hypervariable in length and amino acid composition across FtsZ orthologs (55). We investigated covariations of sequence regions within FtsZ to ask if the interaction between the T7 loop and the CT17 modulated by the CTL is manifest as a statistically significant covariation. For this, we applied the Statistical Coupling Analysis (SCA) first developed by Lockless and Ranganathan. This approach identifies groups of covarying residues, termed sectors, and helps determine the contribution of correlation to positional sequence conservation or covariation (64-66). First, the full sequences of 1208 FtsZ orthologs were collected and aligned using multiple sequence alignment (MSA) (67, 68). The results from the MSA were used for the SCA along with the structure of *Bs*-FtsZ (48) as a reference for residue positions. Then, a covariance matrix between all pairs of sequence positions, weighted by the extent of conservation, was computed (*SI Appendix*, **Figure S3**). Five sectors were identified from the SCA matrix that show correlations across groups. The sectors are arbitrarily named Sector 1 through 5 (**Figure 3B**). As expected, the CTL is not represented in any of the sectors. This implies that it might be under independent selection, distinct from other parts of FtsZ. On the other hand, both the core domain and the CT17 are represented by one of the five sectors.

**Figure 3:**
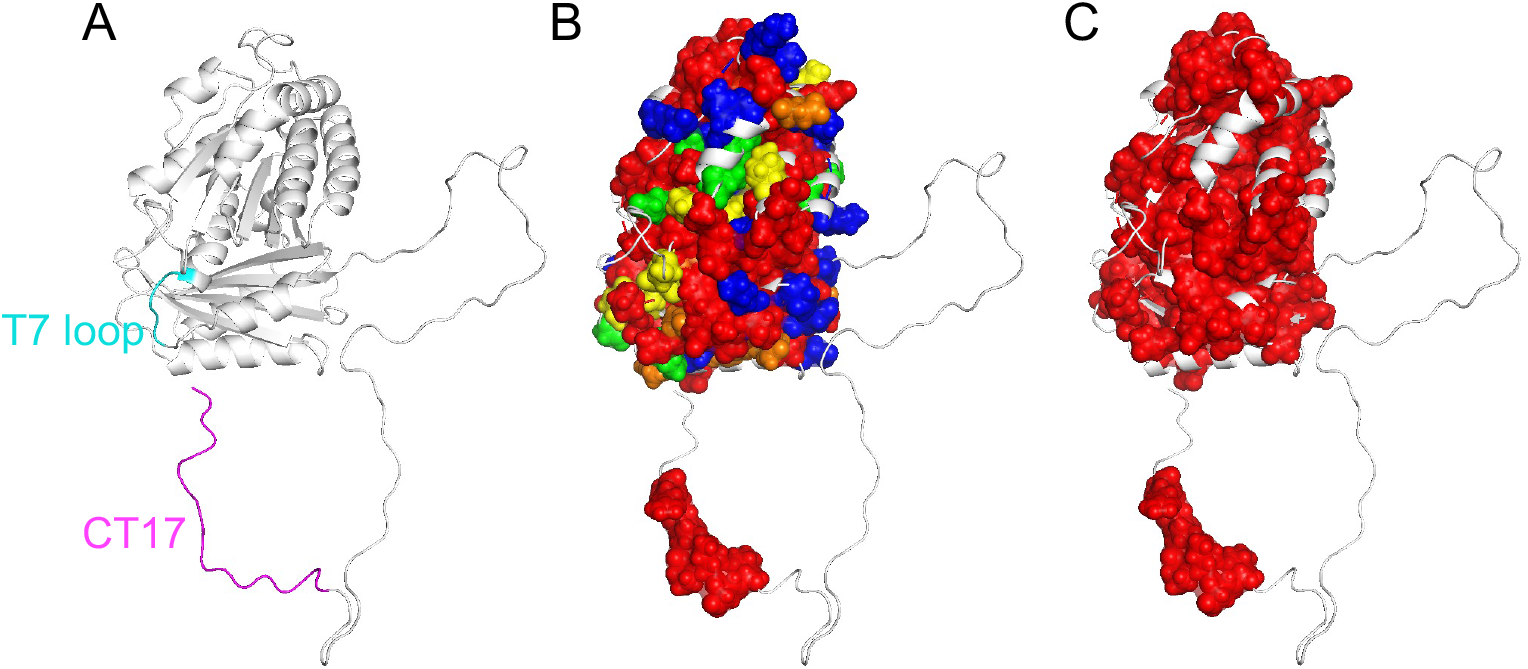
Protein sectors identified within FtsZ show covariation of the CT17 and most of the core domain surface, including the T7 loop. (A) Structure of *Bs*-FtsZ FtsZ is shown (PDB: 2VAM) (48) with cartoon-like schematic of CTT emanating from the C-terminus of the core domain as a reference for panels B and C. The T7 loop is highlighted in cyan and the CT17 in magenta. (B) Five protein sectors are identified from SCA of 1208 FtsZ orthologs. Each of Sectors 1 through 5 is colored in red, orange, yellow, green, and blue, respectively. (C) Only Sector 1 is shown in red, which includes both the T7 loop and the CT17.

Sector 1 is represented by 43% of the core domain residues that are prominently represented on the surface of FtsZ, including a part of the T7 loop (residues 205 and 206), in addition to an 8-residue region in the CT17 (**Figure 3C**). Of particular interest is whether the representation of both the T7 loop and the CT17 in the same sector indicates covariation of the two regions. To examine this, we performed SCA on the sequences of only the core domains that were extracted from the same set of FtsZ orthologs. Interestingly in the core-only analysis, the T7 loop does not belong to any of the sectors, thus appearing as a distinct non-sector module in terms of the core alone (*SI Appendix*, **Figure S4**). However, the nucleotide binding pocket within the core domain, which forms one-half of the dimerization site, remains part of Sector 1 (*SI Appendix*, **Figure S4**). The presence of T7 loop residues and the nucleotide binding pocket in Sector 1 or any of the sectors is only realized in the presence of CT17, indicating covariation of the T7 loop and the CT17 regions. These results suggest that tethering enables covariation of regions that engage in autoinhibitory interactions. The next question is if the CTL, which enables the tethering, is a random, flexible IDR or if there are distinct sequence patterns that facilitate optimal autoinhibitory and other functions.

### Identifying non-random sequence patterns within the Bs-FtsZ CTL

Next, we asked if there are non-random sequence patterns within the CTL of *Bs-*FtsZ. Our goal was to identify sequence patterns that are likely facilitators of CTT interactions with the core across FtsZ orthologs. For this, we utilized the recently described NARDINI algorithm to analyze the CTL of *Bs-*FtsZ and extract non-random binary patterns using this alignment-free approach (69). NARDINI quantifies the extents of mixing vs. segregation of different pairs of amino acid types and ascribes a *z*-score to each of the binary patterns. A pattern is deemed to be non-random if the associated *z*-score meets the criteria of −1.5 < *z*_xy_ or *z*_xy_ > 1.5, where xy refers to a specific pair of residue types. This analysis shows that the CTL of *Bs*-FtsZ is characterized by non-random linear segregation of polar residues x ≡ (Gln, Ser, Thr, Asn, His) with respect to negatively charged residues y ≡ (Asp, Glu), positively charged residues y ≡ (Arg, Lys), and proline residues (**Figure 4A**). Additionally, negatively charged residues also show non-random linear segregation with respect to all other residue types. The non-random linear segregation of polar and negatively charged residues appears to explain the observed statistically significant, albeit weak interactions with the core (**Figure 2**). Clusters of negatively charged residues are likely to repel the acid-rich surface of the core (**Figure 1A**) whereas clusters of polar residues are likely to enable weak interactions with the core. These features of the CTT, specifically the CTL, appear to be facilitators of the observed interactions between the T7 loop and CT17. Interestingly, positively charged residues are randomly distributed along the sequence of the CTL (*z*_++_ < 1.5), indicating a selection against clusters of positively charged residues that can interact favorably with the negatively charged surface of the core (**Figure 1A**).

**Figure 4:**
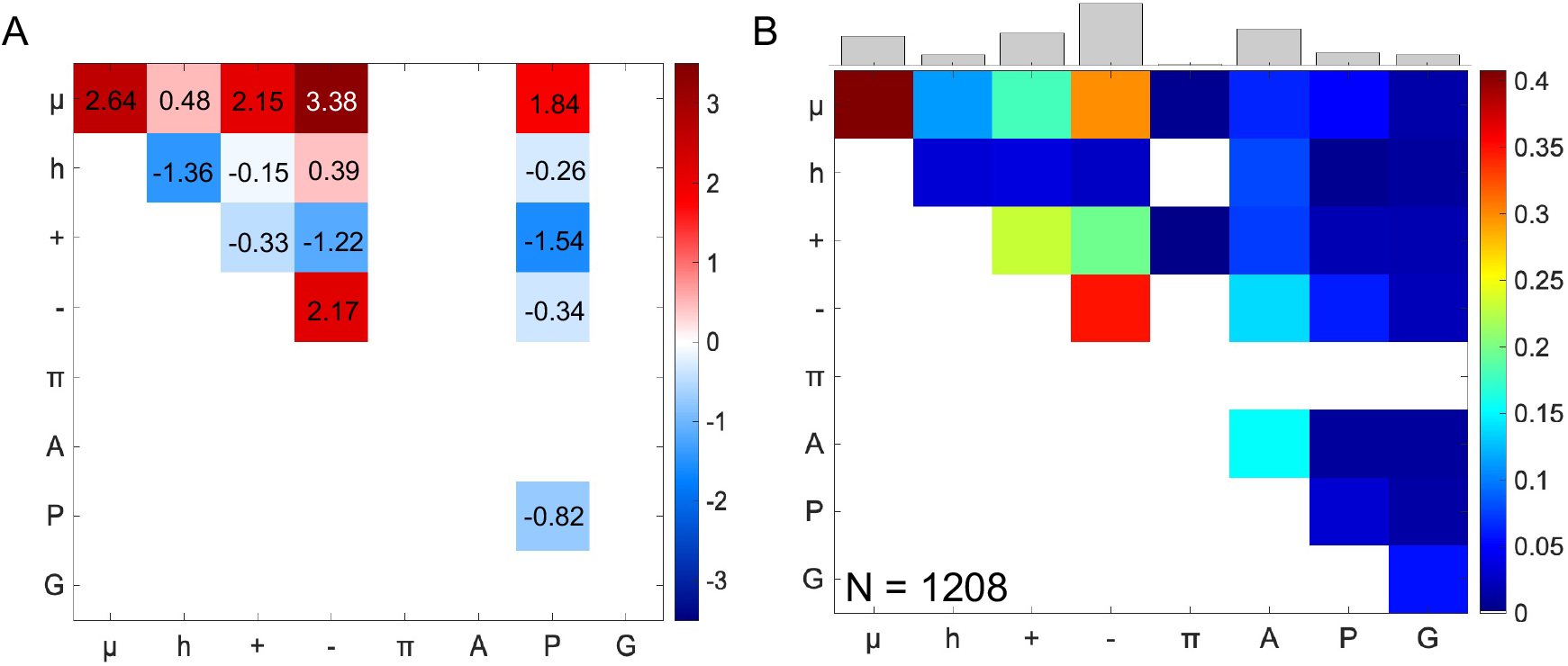
Uncovering non-random binary patterns in the *Bs*-FtsZ CTL and from CTLs across orthologs. (A) Z-score matrix for the CTL of *Bs-*FtsZ shows statistically significant non-random sequence patterns. Color bar indicates the z-scores. The axes describe the residue or residue types for which the z-scores were calculated: μ = polar, h = hydrophobic, + = positively charged, - = negatively charged, π = aromatic, A = alanine, P = proline, G = glycine. (B) Frequency of observing non-random segregation of different types of residue pairs (z_xy_ > 1.5) in CTLs from 1208 FtsZ orthologs. The histogram on top shows relative cumulative frequencies of non-random features for segregation of each residue / residue type.

Despite the demonstrated hypervariability of CTL sequences (55), we find that several binary sequence patterns are conserved across CTLs derived from 1208 distinct FtsZ orthologs (*SI Appendix*, **Figure S5**). In accord with findings for *Bs-*FtsZ, we find that polar and negatively charged residues form linear clusters that are segregated from one another and all other residue types (**Figure 4B**) within CTL sequences. The NARDINI analysis highlights the conservation of linear clusters of negatively charged residues and clusters of polar residues. We propose that these features enable the preservation of statistically significant, albeit weak CTL-core interactions that enable inhibitory interactions through contacts between the CT17 peptide and the T7 loop. The corollary of our proposal is that disruption of the conserved sequence patterns, achieved by mixing or segregating oppositely charged residues along the sequence of the CTL, will weaken native interactions, or enhance non-native interactions, thus having phenotypic consequences *in vitro* and *in vivo*. We tested this hypothesis using sequence design whereby we generated variants, each featuring CTLs with the same amino acid composition of the WT *Bs-*FtsZ while titrating the extent of mixing vs. segregation of oppositely charged residues within the CTL.

### Design of C-terminal linker variants

It is well-known that IDRs where the oppositely charged residues are well-mixed in the linear sequence will prefer expanded, well-solvated conformations (70-72). Additionally, CTLs with well-mixed oppositely charged residues in the linear sequence are expected to weaken native CTT interactions with the negatively charged core. Conversely, IDRs where the oppositely charged residues are segregated with respect to one another have the potential to engage in strong non-native intra-CTL / intra-CTT interactions, non-native interactions with the FtsZ core, and non-native inter-CTL interactions between different FtsZ molecules.

The linear mixing / segregation of oppositely charged residues can be quantified using different parameters (70, 73, 74) including κ_+-_, where 0 ≤; κ_+-_ ≤; 1 (70). Oppositely charged residues that are uniformly mixed within a linear sequence will yield values of κ_+-_ ≈ 0 whereas a separation of oppositely charged residues into discrete blocks will yield values of κ_+-_ ≈ 1. To test the impact of the linear segregation vs. mixing of oppositely charged residues on FtsZ functionalities, we designed a set of CTL variants using the amino acid composition of the CTL from *Bs-*FtsZ. The designed sequences span κ_+-_ values between 0.14 and 0.72 (*SI Appendix*, **Table S1**). The calculation of CTT κ_+-_ values includes both the CTL and CT17 modules. Each of the designed *Bs-* FtsZ variants are denoted as K*x*, where *x* is hundred times the κ_+-_ value of the CTT in question. The sequences of the core domain and the CT17 were not altered.

### Increasing κ_+-_ of the CTT introduces intra-CTT interactions and enhances CTT-core interactions

The working hypothesis is that increased charge segregation enhance non-native intra-CTT thereby weakening CTT-core interactions and reducing the potential for autoinhibition. To test this hypothesis, we performed two types of simulations of the Kappa variants of *Bs-*FtsZ. In the first mode, we performed simulations of CTT variants as autonomous units. In the second mode, we interrogated the conformations accessible to full-length FtsZ molecules with the CTT tethered N-terminally to the core. Results from simulations in mode 1 show a decrease of ensemble-averaged radii of gyration (*R*_g_) as κ_+-_ is increased (**Figure 5A**). This derives from the increased favorability of intra-CTT interactions between blocks of oppositely charged residues within the CTL as the segregation of oppositely charged residues is increased. We tested predictions from simulations by performing fluorescence correlation spectroscopy (FCS) measurements to quantify the ensemble-averaged hydrodynamic radii (*R*_h_) of CTT peptides. We observe a clear one-to-one correspondence between the calculated *R*_g_ and measured *R*_h_ values. It follows, in accord with previous observations (70, 71, 75), that increasing the linear segregation of oppositely charged residues drives the compaction of IDRs through cohesive interactions within the IDRs.

**Figure 5:**
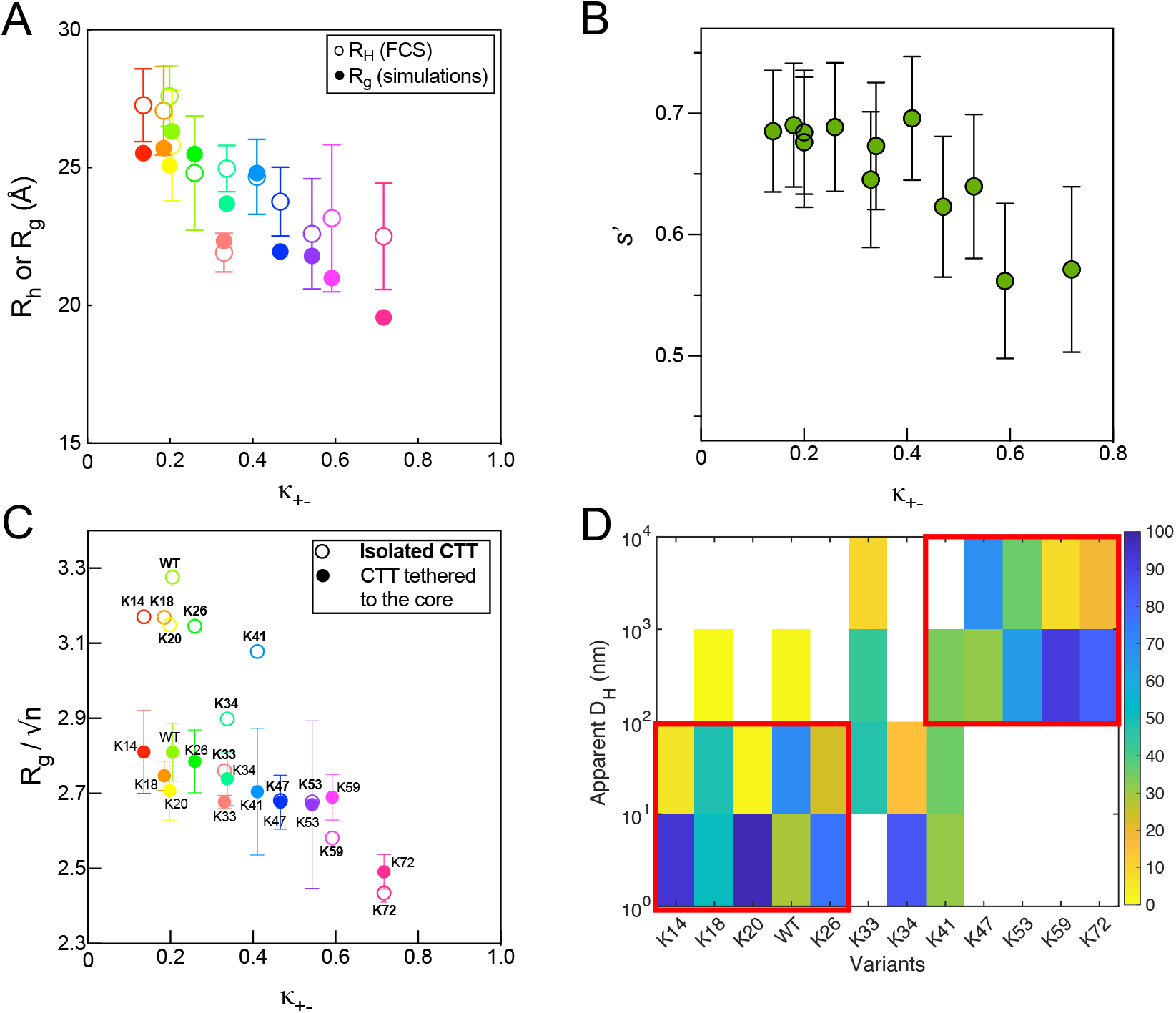
Increased linear segregation of oppositely charged residues leads to compaction. (A) Computed *R*_g_ values from simulations and measured *R*_h_ values from FCS experiments as a function of κ_+-_ values of the Kappa variants. (B) Calculated Shannon entropy (*s*′) values of the Kappa variants from computed bivariate distributions of *R*_g_ and asphericity (δ*) values. (C) Normalized *R*_g_ of the Kappa variants from simulations performed in two modes, isolated (open symbols) and tethered to the GTPase core (solid symbols). CTT sequences with lower κ_+-_ values undergo compaction upon being tethered to the folded core. (D) DLS data, shown as a checkerboard plot, quantifies the *percent* likelihood of observing a scatterer of a specific size for each of the Kappa variants.

Two-dimensional histograms of normalized *R*_g_ and asphericity values of the CTT variants show evidence of conformational heterogeneity as manifest in the broad distribution of the sizes and shapes (55) (*SI Appendix*, **Figure S6**). We used these distributions to compute information theoretic measures of conformational heterogeneity such as relative Shannon entropies (*s*′) (55) (**Figure 5B**). Despite a doubling of κ_+-_ from 0.20 and 0.41, the values of *s*′ for the relevant CTT variants are bounded between 0.65 and 0.7. This implies that despite the decrease in *R*_g_ with increasing κ_+-_, the overall conformational heterogeneity is not greatly impacted because enhancements in shape fluctuations offset any diminution in size fluctuations (55). However, we observe a clear reduction in *s*′ values as κ_+-_ increases beyond 0.41.

Simulations of full-length proteins with the CTTs attached N-terminally to the core domain (**Figure 5C**) show that the CTT undergoes additional compaction for κ_+-_ < 0.47. This derives from contacts formed between the CTT and the core domain. However, this additional compaction is not observed in variants where the CTT κ_+-_ is greater than 0.41. This is because the interactions between oppositely charged blocky residues within the CTT are stronger than CTT-core interactions as κ_+-_ increases beyond 0.41. The simulations support the hypothesis that segregation of oppositely charged residues into distinct blocks strengthens intra-CTT interactions thereby minimizing CTT-core interactions. This should weaken the autoinhibitory functions of the CTT, leading instead to alternative interactions in *cis* and in *trans* via the CTT.

### CTT-CTT interactions can drive non-native FtsZ assemblies in vitro

The preceding discussion shows that linear segregation of oppositely charged residues within the CTL introduces non-native interactions within the CTL. We reasoned that increased segregation of oppositely charged residues within the CTL has the potential of introducing non-native, CTT-mediated inter-domain interactions in *trans*. These are likely to involve a combination of CTT-core and CTT-CTT interactions in *trans*, that are weak or non-existent for the wild-type FtsZ. To probe for these non-native interactions, we used dynamic light scattering (DLS) measurements of the different *Bs-* FtsZ variants to assess the contributions of non-native intermolecular interactions between FtsZ protomers. These measurements were performed in the absence of GTP to detect the presence of non-native interactions in the designed variants. We observed two groups of apparent hydrodynamic diameters (D_h_) (**Figure 5D**). Diameters smaller than 10^2^ nm correspond to variants whose CTT κ_+-_ values are lower than 0.3. In contrast, D_h_ values larger than 10^2^ nm were observed for *Bs-*FtsZ variants with CTT κ_+-_ values that are greater than 0.4. This increase in assembly size indicates a gain of non-native interactions arising from altered sequence patterns within the CTL.

The contributions of non-native CTT-CTT and CTT-core interactions to the overall FtsZ assembly in the absence of GTP can be inferred by quantifying correlations between D_h_ and the effective interaction strengths of CTT-CTT and CTT-core interactions. The weighted mean of the apparent D_h_ from DLS measurements was used as a quantitative proxy for the strengths of driving forces for *Bs-*FtsZ assembly in the absence of GTP. The effective strengths of CTT-CTT and the CTT-core interactions were extracted from statistical analysis of simulation results. First, the effective strengths ε_inter-CTT_ of CTT-CTT interactions were determined by extracting *R*_g_ values using snapshots where the CTT is not in contact with the core. This helps us examine how strongly the CTT interacts with itself. The quantity of interest was defined as: 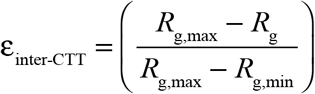 where *R*_g_ is the construct specific mean radius of gyration and *R*_g,max_ and *R*_g,min_ are the maximum and minimum values of the radii of gyration across κ_+-_ variants, respectively (**Figure 6A**). The derived values of ε_inter-CTT_ are positively correlated with D_h_ (linear regression Pearson *R*^2^ = 0.62) indicating that inter-CTT interactions contribute to the FtsZ assembly in the absence of GTP. Second, the effective strengths of CTT-core interactions were calculated over snapshots where the CTT is in contact with the core using *p*_c_, the mean per residue probability of finding any CTT residue in contact (5 Å) with any residue on the core in the simulated ensemble. Thus, *p*_c_ is large if many CTT residues interact with the core and the parameter of interest is defined as: 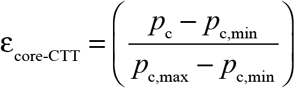. Here, *p*_c,max_ and *p*_c,min_ are the maximum and minimum values of *p*_c_ across Kappa variants. The correlation between ε_core-CTT_ and D_h_ is poor (linear regression Pearson *R*^2^ = 0.21). This implies that CTT-core interactions on their own make negligible contributions to the driving forces for FtsZ assembly in the absence of GTP (**Figure 6B**).

**Figure 6:**
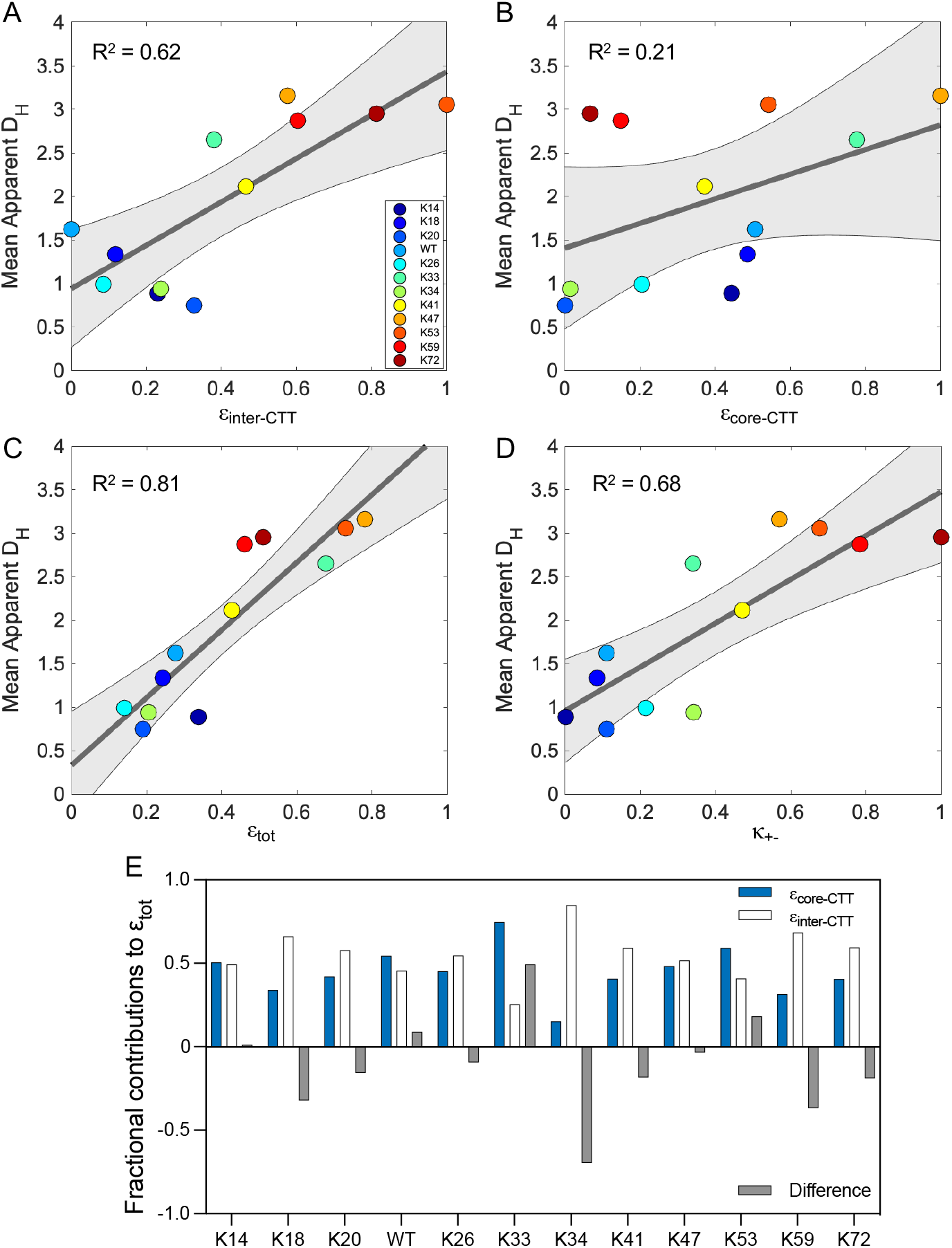
CTT-CTT and CTT-core interactions contribute to FtsZ assembly in the absence of GTP. Correlation between the mean apparent D_h_ measured using DLS and (A) ε_intra-CTT_ (B) ε_core-CTT_ (C) ε_tot_ and (D) κ_+-_. Here, the grey area is the 95% confidence region of the linear regression. (E) Fractional contributions of ε_inter-CTT_ (white bars) and ε_core-CTT_ (blue bars) to ε_tot_.

Next, we calculated the coupled contributions of the two parameters to GTP-independent assembly of FtsZ variants. For this, we used a mixture model: ε_tot_ = *f* ε_inter-CTT_ ± (1− *f*)ε_core-CTT_, where 0 ≤; *f* ≤; 1. Here, *f* is the fraction of conformations in which the CTT is not in contact with the core. Joint consideration of the contributions of both types of interactions yields a strong positive correlation between ε_tot_ and D_h_ (linear regression, *R*^2^ = 0.81) (**Figure 6C**) providing we extract a unique, Kappa-variant specific value of *f* (**Figure 6E**). The correlation between D_h_ and κ_+-_ is weaker than that between D_h_ and the total effective drive to assemble (linear regression Pearson *R*^2^ = 0.68, **Figure 6D**). This is because κ_+-_ only describes the CTT-CTT interaction strength according to the positioning of oppositely charged residues, whereas the mixture model accounts for the contributions of CTT-core interactions. Taken together, our results point to a combination of variant-specific CTT-CTT and CTT-core interactions as contributors to non-native FtsZ assembly in the absence of GTP. CTT-core interactions within the Kappa variants are not identical even if the values of *f* are similar. For example, *f* ∼ 0.5 for K14, WT, K26, and K47, and yet the interaction between the CT17 and the T7 loop is abolished in K47 (*SI Appendix*, **Figure S7C, S7D**). Instead, we observe a gain of non-native CTL interactions with the T7 loop in this variant (**Figure S7A, S7B**). Overall, the pattern of non-native interactions will be influenced by the totality of non-random sequence patterns within the designed CTLs (**Figure 4**, *SI Appendix*, **Figure S7**).

### Non-native interactions driven by redesigned CTLs promote large and diverse assemblies that are less active in GTP hydrolysis than WT

We investigated the effects of altered sequence patterns in the Kappa variants on GTP-dependent assembly and GTP hydrolysis. *In vitro, Bs-*FtsZ forms single-stranded protofilaments that associate laterally to form bundles (17, 49, 76-79). Right-angle light scattering is a sensitive method for studying FtsZ polymerization and bundling of FtsZ polymers in the presence of GTP (56). We measured scattering intensities of the Kappa variants and normalized these to measured values of WT for comparison. We observed two broad categories of variants using a 3-fold increase in scattering intensities when compared to WT as a threshold (**Figure 7A**). In general, the variants where the CTT κ_+-_ is ≤; 0.41 show lower scattering intensities and those with the CTT κ_+-_ values greater than 0.41 show higher scattering intensities. However, K18 is an exception. Despite having a CTT κ_+-_ value that is close to that of WT, the K18 variant exhibits high scattering intensities, indicating the formation of large assemblies.

**Figure 7:**
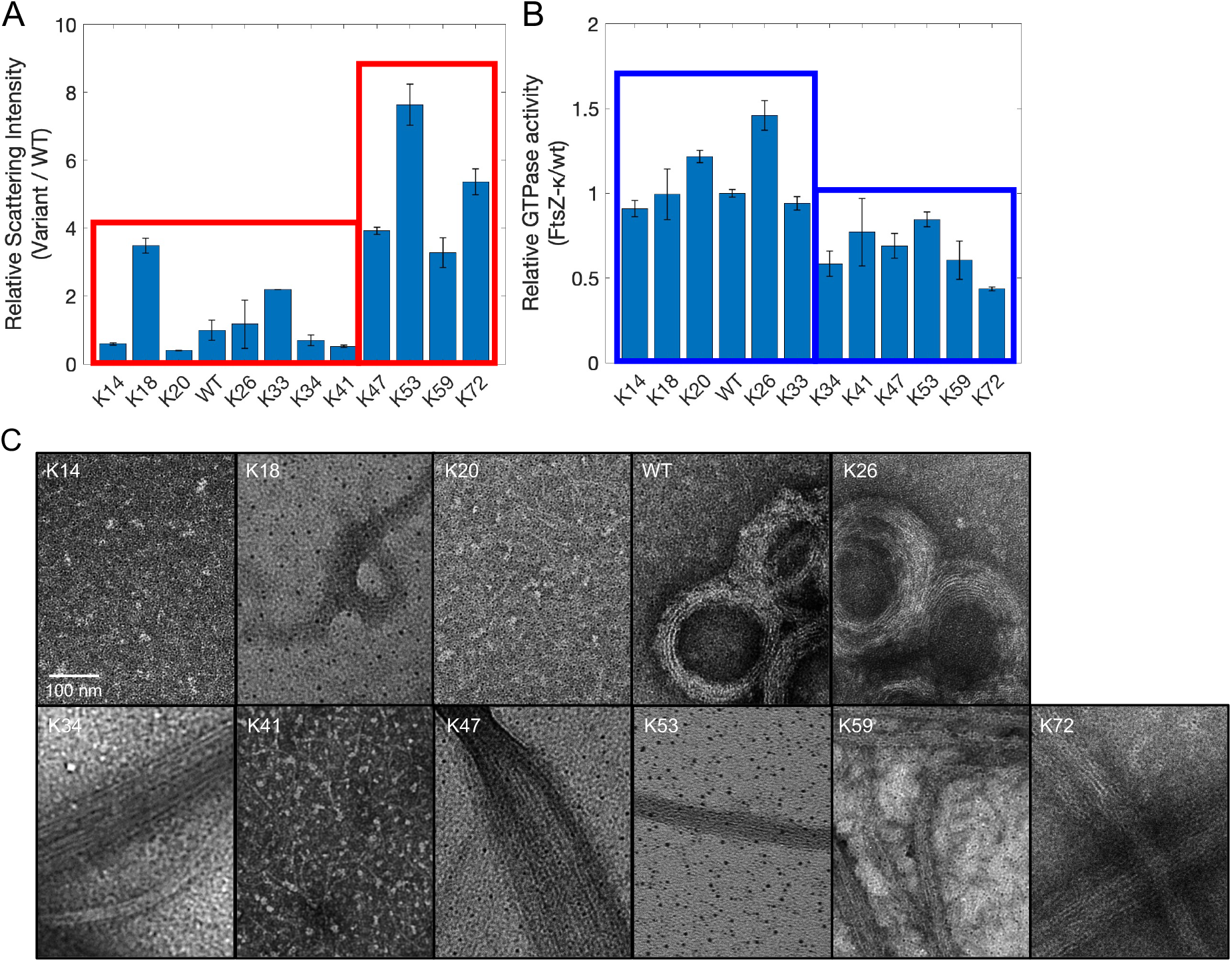
Changes to the linear mixing vs. segregation of oppositely charged residues within the CTL affects GTP-dependent FtsZ assembly and GTPase activity. (A) Scattering intensities of 5 μM of WT FtsZ and different variants were measured in the presence of 1 mM GTP. Values reported here were normalized to those of WT. The red outlines indicate two broad categories of the Kappa variants that show less than or more than 3-fold increase in scattering intensities compared to WT. (B) Rates of GTP hydrolysis of the Kappa variants are normalized to that of WT. Blue outlines indicate two broad categories of the Kappa variants that show faster, or slower rates compared to the WT. (C) Negative-stain TEM micrographs show the diverse morphologies formed by the Kappa variants.

Negative-stain transmission electron microscopy (TEM) imaging helps capture the diverse morphologies of higher-order assemblies formed by the Kappa variants (**Figure 7C**). Those that show lower scattering intensities correspond to protofilaments – see data for K14, K20, K33 (56), K41 – or rings composed of laterally associating bundled protofilaments (WT, K26). On the other hand, stronger non-native interactions driven by segregation of oppositely charged residues within the CTL lead to increased self-association of FtsZ variants via inter-CTT interactions among filaments. These interactions generate profoundly different assemblies when compared to morphologies observed for WT *Bs-*FtsZ (17, 30, 31, 80-82). Increase in the CTT κ_+-_ leads to the formation of long, linear filamentous tracts characterized by inter-filament interactions that appear to involve the CTTs – see data for K47, K53, K59, and K72 in **Figure 7C** and *SI Appendix*, **Figure S8**. These long FtsZ polymers are likely to compromise treadmilling thereby suppressing the exchange of FtsZ subunits in polymers upon GTP hydrolysis.

Next, we measured the rates of GTP hydrolysis of Kappa variants using a series of independent continuous assays for GTPase activity. Each measurement was performed with 5 μM FtsZ in buffer MES with 1 mM GTP. As with the right-angle scattering results, the Kappa variants can be grouped into two broad categories based on their GTPase activity relative to WT (**Figure 7B**). The concentration-specific GTPase activities are likely to be a convolution of many variables, including multiple assembly states of FtsZ and CTL-dependent variations to Michaelis-Menten parameters. However, at identical concentrations of FtsZ subunits, the sequence-encoded features within different CTLs directly influence the functions of the GTPase core domain with extreme segregation of oppositely charged residues reducing the GTPase activity.

Taken together, our results suggest that the sequence patterning, and the emergent native contacts lost, or non-native contacts that are formed in *cis* (intra-CTL and CTT-core) and *trans* (CTT-CTT and CTT-core) can alter the functions of *Bs-*FtsZ variants. We find that extreme segregation or mixing of oppositely charged residues in the CTL cause significant deviations from WT-like behavior in FtsZ assembly and enzymatic activity *in vitro*. However, we also observe robustness of both functions for CTT κ_+-_ values between 0.19 and 0.34.

### Extremes of mixing or segregation of oppositely charged residues show pronounced in vivo phenotypes

In contrast to the assayed *in vitro* functionalities, *in vivo* functions of FtsZ are reliant on factors such as protein stability in the cell and interactions with other components of the division machinery. To evaluate *in vivo* functionalities of the Kappa variants, we used a strain that would allow us to examine variants as the sole copy of FtsZ regardless of their ability to support cell growth and division. This strain was made by cloning variant alleles into a *B. subtilis* strain where the sole WT *ftsZ* gene is under the control of a xylose inducible promoter and the variant allele is under the control of an independent Isopropyl β-d-1-thiogalactopyranoside (IPTG) inducible promoter (42). We can then deplete WT FtsZ and induce the expression of the variants in a controlled manner.

We first evaluated the stability of FtsZ in cells using quantitative immunoblots since we observed notable changes in conformational ensembles with changes in CTT κ_+-_ values (**Figure 5C and** *SI Appendix*, **Figure S6**), and it is reasonable to expect that changes to protein structure can lead to degradation *in vivo*. Indeed, the ClpX chaperone is reported to modulate assembly of FtsZ in *B. subtilis*, and the altered conformations of CTT may lead to increased susceptibility for ClpX-mediated proteolysis by ClpP (83, 84). Furthermore, the ClpXP complex is reported to interact with the disordered CTT of FtsZ to enhance degradation in *E. coli* (85). Samples were prepared from mid-exponential phase cultures (OD600 of ∼0.5) that were back diluted into IPTG, then grown to and harvested at mid-exponential phase. Immunoblots were normalized to total protein levels. All but three variants accumulated near or above WT levels. The unstable variants are those with CTTs featuring extreme κ_+-_ values (**Figure 8A**) *viz*., K14, K53, and K72. These variants appear to be substantially degraded (< 20% WT FtsZ levels), indicating that they are more susceptible to proteolysis.

**Figure 8:**
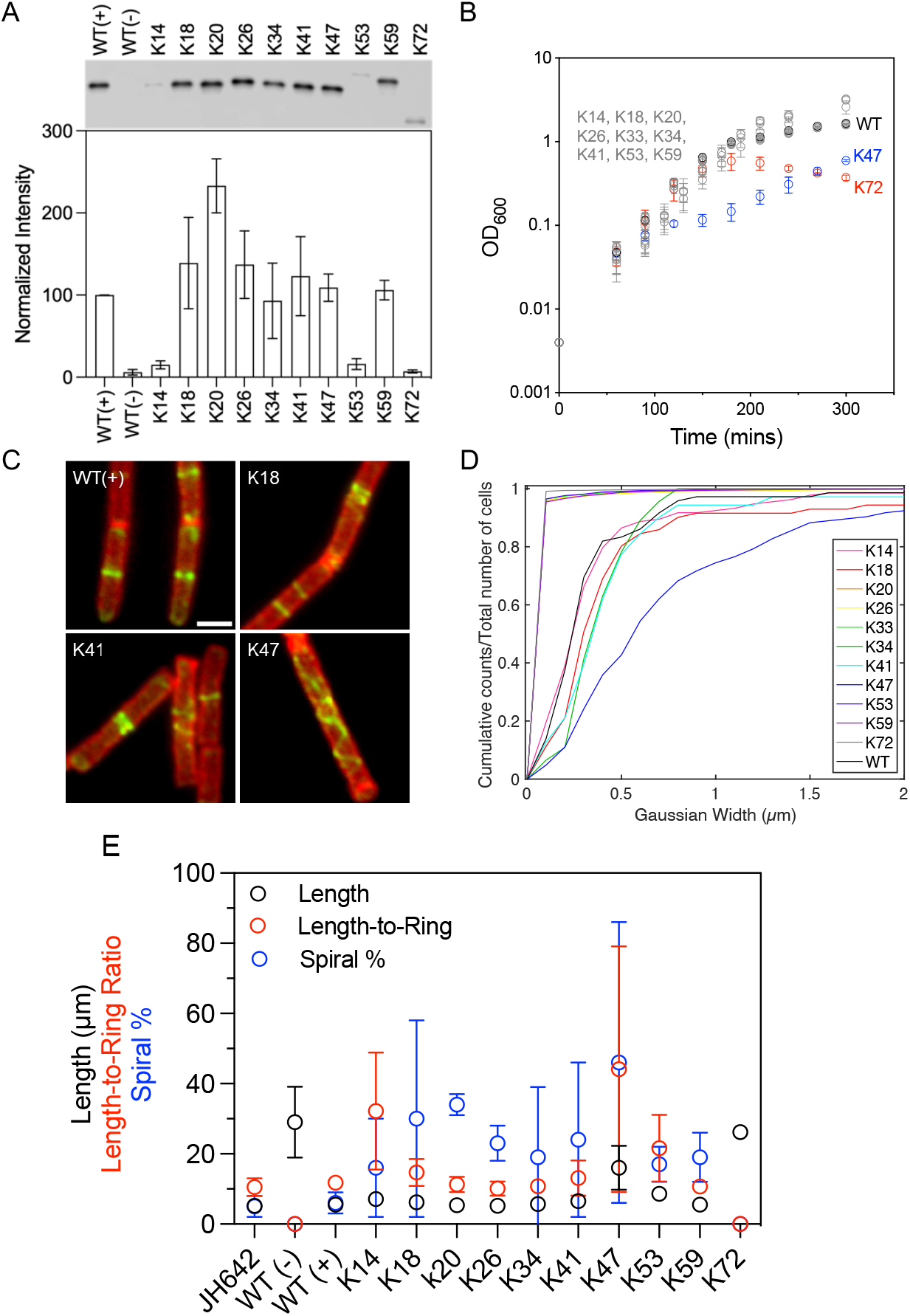
Changes to the extent of segregation vs. mixing of oppositely charged residues within the CTL affects protein stability, cell growth, and division. (A) Quantitative immunoblots of the Kappa variants. Intensities from the gel (top) are quantified in the histogram (below) and normalized to the WT. (B) Cell growth profiles of the Kappa variants. (C) In the IFM images, FtsZ is artificially colored in green, and the cell wall is in red. Scale bar = 2 μm. (D) The extent of condensation of FtsZ Kappa variants in cells is quantified using cumulative distribution functions. (E) We quantified cell lengths, length-to-ring ratios, and percentage of cells with spirals from the IFM images. Data for at least 200 cells were used for each of the Kappa variants. The L/R data for WT(-) and K72 were plotted at 0 since neither rings nor spirals were observed.

Consistent with poor stability, strains expressing K72 were unable to support growth of *B. subtilis* (**Figure 8B**). In a clear contrast, the control strain (PAL3171) expressing only WT *ftsZ* under the IPTG inducible promotor exhibited a standard growth curve with exponential and stationary phases. As expected, the same control strain without *ftsZ* expression grew for approximately three hours before plateauing and slightly decreasing in optical density (86). Interestingly, all Kappa variants except for K47 and K72 exhibited robust WT-like growth profiles despite degradation of K14 and K53 in cells. Strains expressing K47 were able to support growth, but grew more slowly than the WT despite having FtsZ levels equivalent to WT. This suggests that K47 may interfere with both cross-wall synthesis and synthesis of the longitudinal cell wall that leads to defects in growth.

To gain insight into FtsZ assembly *in vivo*, we used immunofluorescence microscopy (IFM) to quantify cell length and image FtsZ ring formation. Cultures were prepared as described for immunoblots, but harvested after roughly four doublings (OD600 of 1.5-0.3) for fixation and imaging (49). Longer cell lengths indicate impaired cell division. Cells expressing the three unstable variants, K14, K53, and K72, were 30%, 56% and 376% longer than cells expressing WT (**Figure 8E**, *SI Appendix*, **Figure S9**). Despite the high κ_+-_ value of its CTT, the average length of cells expressing K59 was akin to that of WT, while cells expressing the K18 allele were ∼13% longer than WT cells. The variants with extreme segregation or mixing of the oppositely charged residues in their CTTs that exhibited longer cell lengths, also exhibited poor Z-ring formation. The Length-to-Ring ratio (L/R) metric was used to evaluate the ability of FtsZ to form Z-rings, and is defined as the ratio between the cumulative length of all measured cells and the cumulative numbers of rings in those cells (87). K14 and K47 showed higher L/R ratios, K72 was unable to form any rings, whereas K59, despite the higher κ_+-_ value of the CTT showed robust WT-like protein stability, cell growth profile, cell length, and L/R.

Images of cells expressing Kappa variants revealed the presence of unusual FtsZ localization patterns in many of the strains (**Figure 8C**). WT FtsZ forms well-defined medial rings with residual localization at cell poles. In contrast, most variants displayed loose, spiral-like structures in addition to distinct rings. Previous work identified spirals as being indicative of defects in lateral interactions between single stranded polymers which interferes with condensation into precisely localized Z-rings (88-91). Accordingly, we quantified the spiral phenotype by quantifying the percentage of cells with spirals. This quantity, which we termed Spiral %, was higher in all variants when compared to cells expressing only WT *Bs-*FtsZ. We also computed, the Gaussian width (**Figure 8D**), which is a measure of spiral width relative to cell length. Spiral formation was independent of FtsZ levels, cell length, and L/R ratios. Only K72 lacked spirals possibly due to having insufficient levels of FtsZ for assembly (**Figure 8A**). This suggests that the Kappa variants are unable to condense efficiently. Instead, deviation from WT-like sequence patterns leads to a mixture of rings and spiral structures (*SI Appendix*, **Figure S10)**. The latter are likely to be intermediate states that represent CTL-driven defects in condensation and / or localization.

## Discussion

Considerable attention has focused on the lengths of CTLs and their impacts on FtsZ functions (82, 92-94). Conclusions from such studies can be confounding because the CTLs are not simple homopolymers. As a result, titrating lengths in arbitrary ways, creates a multivariable problem where the composition and sequence patterning of residues are also varied, thereby confounding the inferred relationships between length and function. To avoid such problems, we focused on a single sequence parameter κ_+-_ whereby we fixed the length and amino acid composition and directly modulated the interplay between intra- and intermolecular interactions. Our results show that changes to sequence patterns within the CTL have clear biophysical, biochemical, and biological consequences. We generally observe deleterious consequences in FtsZ functionalities when the oppositely charged residues of the CTL are segregated with CTT κ_+-_ values greater than 0.34 leading to aberrant assemblies. This gain-of-function affects both the CTT-core and CTT-CTT interactions between FtsZ molecules. Our results suggest that the loss of native interactions between the T7 loop and the CT17 can be detrimental, as seen for K47. In contrast, the preservation of native contacts in K59 helps maintain WT-like functions *in vivo* because the cognate autoinhibitory interactions are maintained for these variants. Thus, any sequence solution that mediates the native interaction between the T7 loop and the CT17 is a viable CTL. Here, we find at least one sequence solution that does so by non-random segregation of negatively charged residues. Other solutions may exist for sequences that accommodate the conserved non-random sequence patterns.

Taken together, our results suggest that the drive to limit interactions between the CTL and the core is conserved in FtsZ orthologs. This is achieved via non-random segregation of negatively charged residues, and dispersion of positively charged residues along the CTL sequences. These features contribute to the minimization of associations that lead to aberrant multivalent interactions amongst segregated blocks of charge within CTTs, and between the CTT and core. Although we focused on titrating the linear segregation / mixing of oppositely charged residues, one can imagine a generalization of our approach that uses sequence libraries or directed evolution to query how large- and small-scale changes to the assortment of conserved sequence patterns alter the functionalities IDRs. Such an approach was deployed in recent studies to identify rules underlying the functions of disordered transactivation domains of transcription factors (72, 95, 96). Investigations of additional sequence variants that impact the conformational ensemble via alternative CTL-mediated interactions such as those mediated by polar or polyglutamine tracts (97, 98) will be helpful to assess the impact of different types of polar interactions.

Our findings indicate that along with composition and length, an optimal CTL requires the balancing of strengths of several parameters including inter-module interactions in *cis* and in *trans*, the non-random / random sequence patterns, and conserved sequence-ensemble relationships (55). Indeed, the CTT / CTL encoded effects may need to be just right – a Goldilocks effect – given that the CTT and its two distinct modules play multiple regulatory roles. Similar results have been uncovered in a recent study of hypervariable linkers within viruses (99).

## Materials and Methods

All *B. subtilis* strains were derived from the strain JH642 (100). Cloning and genetic manipulation were performed using standard techniques (101, 102). All cloning was done using the *E. coli* strain AG1111 derivative PL930 (103). PL930 contains the low copy plasmid pBS58 expressing *E. coli* ftsQAZ, which facilitates cloning of *B. subtilis* FtsZ. Vent DNA polymerase was used for PCR (New England Biolabs). All restriction enzymes and T4 DNA ligase were purchased from New England Biolabs. All genomic DNA extractions were performed using the Wizard Genomic DNA Purification Kit (Promega). Plasmid preparations were made using the NucleoSpin Plasmid Kit (Macherey-Negel). Gel/PCR purifications were performed using the NucleoSpin Gel and PCR Clean-up Kit (Macherey-Negel). Cells were grown in Luria-Bertani (LB) medium at 37 ºC unless otherwise noted. Antibiotics were used at the following concentrations: ampicillin = 100 μg ml^-1^, spectinomycin = 100 μg ml^-1^, chloramphenicol = 5 μg ml^-1^.

Additional details regarding cloning, strain construction, growth conditions, immunoblotting, measurements of growth curves, IFM, sequence analysis, and details of the simulation setup are described in the *SI Appendix*.

## Acknowledgments

We thank Stephen Vadia and Catherine Kornacki for cloning and strain constructions. TEM imaging was performed in part using the Washington University Center for Cellular Imaging (WUCCI) supported by Washington University School of Medicine, The Children’s Discovery Institute of Washington University and St. Louis Children’s Hospital (CDI-CORE-2015-505 and CDI-CORE-2019-813) and the Foundation for Barnes-Jewish Hospital (3770 and 4642). MKS was a CSELS postdoctoral fellow of the erstwhile Center for Science & Engineering of Living Systems (CSELS) in the James McKelvey School of Engineering at Washington University in St. Louis. This work was also supported by the Air Force Office of Scientific Research (FA9550-20-1-0241) and the US National Science Foundation (MCB1614766) to RVP, and 5R35GM127331 from the US National Institutes of Health to PAL.

