## Supporting Information Appendix for "Connecting sequence features within the disordered C-terminal linker of *B. subtilis* FtsZ to functions and bacterial cell division"

for

### Supporting Information Text

**Cloning:** Synthetic oligonucleotides of the Kappa variants were synthesized (Integrated DNA Technologies) and PCR amplified, restriction digested, and inserted into pPJ19, which contains FtsZ under the control of the $P_{spac}$ promoter that is inducible with IPTG, with restriction sites flanking the CTL. A BamHI site after residue 315 and an XmaI site before residue 366 result in the insertion of amino acid pairs GS and PG N- and C-terminal to the CTL, respectively. All DNA sequences were confirmed by Sanger sequencing.

**Strain construction:** The Kappa variant strains were constructed as described previously (1, 2). Strains and plasmids used in this study are listed in *SI Appendix*, **Table S1.** They were constructed by first transforming plasmids into wild-type JH642 cells. This strain permits depletion of wild-type FtsZ and is xylose-dependent for normal growth (3).

**Growth conditions:** *Bacillus subtilis* strains were first grown from a single colony overnight in lysogeny broth (LB) containing spectinomycin and xylose to a final concentration of 0.5%. Cells were grown overnight in LB medium at 37°C with 100 µg/mL spectinomycin and 0.5% xylose. They were then diluted 1:100 and grown in fresh LB medium supplemented with 0.5% xylose until $OD_{600}$ reached 0.4-0.5. Cells were then washed twice with LB, diluted 1:100, and grown to mid-log phase in 0.1 mM IPTG.

**Immunoblotting:** Immunoblotting was performed as described previously (7). The cells were lysed with lysozyme and a Fastprep-24 benchtop homogenizer (MP Biomedicals) with 01.mm silica beads. Loading was normalized to the OD600 after pellet resuspension. After transfer to the blot, total protein was quantified using the Spyro Ruby protein blot stain (Thermo Fisher) and imaged on a Gel Dock EZ Imager (Bio-Rad). The blot was probed using affinity-purified polyclonal rabbit anti-FtsZ antibodies and goat anti-rabbit antibodies conjugated to horseradish peroxidase (Invitrogen). Immunoblots were developed using the ECL Western Blotting detection reagents (Bio-Rad Laboratories) and visualized with the luminescent image analyzer Odyssey Fc Imaging System (LI-COR Biosciences).

**Growth Curves:** Cells were grown overnight under the same media conditions as the immunoblots in 0.5% xylose, back diluted 1:100, and grown in 0.5% xylose until the cells reached the mid-log phase. The cells were then washed twice with LB, diluted 1:100, and grown in 0.1 mM IPTG. Starting 1 hour after induction, the $OD_{600}$ was measured approximately every 30 minutes for 7 hours, with later time points taken further apart.

**Immunofluorescence microscopy (IFM):** IFM was performed as described previously (2). Cells were grown using the same media conditions overnight in 0.5% xylose, back diluted 1:100, and grown in 0.5% xylose until the cells reached the mid-log phase. The cells were then washed twice with LB, diluted to OD600 0.025, and grown in 0.1 mM IPTG for five generations (~2.5 hours). The cells were harvested and fixed with 16% paraformaldehyde/0.5% glutaraldehyde. The cells were lysed with 2 mg/mL lysozyme. FtsZ was detected with affinity-purified polyclonal rabbit anti-FtsZ serum combined with goat anti-rabbit serum conjugated to Alexa488 (Life Technologies). Cell walls were stained with wheatgerm agglutinin conjugated to tetramethylrhodamine, and DNA was stained with DAPI. Slides were visualized with an Olympus BX51 microscope with Chroma filters and a Hamamatsu OrcaERG camera. Image capture was performed using Nikon Elements (Nikon Instruments) and analyzed via ImageJ (4). The cell length/Z-ring (L/R) ratio was calculated as previously described (5). The L/R ratio was calculated as the sum of the total cell length of a population of cells divided by the total number of Z-rings in that population. Percent Spirals was calculated as a percent of the sum of all Z rings divided by the total number of FtsZ structures in that population.

Line profiles were drawn across each cell using MicrobeJ plug-in (6) in for ImageJ (4). The intensity profiles were imported and adjusted to subtract background signals. The adjusted profiles were each fitted to a Gaussian using a custom script in MATLAB using: $I(x) = a\exp\left[-\left(\frac{x-b}{c}\right)^2\right]$ where $I(x)$ is the intensity at position $x$. The fits were constrained to yield positive values of $a$, $b$, and $c$ and ensure that $b$ and $c$ were always greater than the cell length. Values of $b$ were binned at 0.1 μm to construct a histogram, from which values were extracted to plot the cumulative distributive functions (CDF) for each of the Kappa variants.

**Protein expression and purification:** Kappa variants were cloned into the pET-21b(+) expression vector through *E. coli* strain AG1111. The resulting plasmids were mini-prepped and freshly transformed into C41(DE3) cells and used for protein expression. 500 mL of LB medium was inoculated 1:100 with an overnight culture. Cells were grown at 37 ºC until $A_{600}$ ~0.6, and then the cells were induced with IPTG to a final concentration of 1 mM. Cells were grown for an additional 4 h at 37 ºC, and then cells were harvested by centrifugation, and cell pellets were stored at -80 ºC. Purification was performed as previously described (2, 7). Protein preparations were concentrated using spin concentrators, separated into aliquots, flash frozen on liquid $N_2$, and stored at –80 ºC. Prior to use in an assay, FtsZ aliquots were thawed on ice and centrifuged to remove any insoluble polymers. FtsZ concentration was then determined using Pierce 660 nm assay (Thermo Fisher Scientific) with porcine tubulin (Cytoskeleton) as a standard.

**90° Light Scattering Assays:** Assembly reactions were performed with 5 μM FtsZ in Buffer MES (50 mM MES, 50 mM KCl, 2.5 mM $MgCl_2$, 1 mM EGTA, pH 6.5) at 30 ºC. Measurements were recorded every quarter of a second. A 1-minute baseline was established before adding 1 mM GTP to the reaction. At least three trials were conducted for each variant. All data were collected and exported into Microsoft Excel, and the subsequent analysis was performed using MATLAB. The average baseline was subtracted from each data point.

**Transmission Electron Microscopy (TEM):** Samples were prepared in conditions mimicking the light scattering assays, with a lower concentration of 2.5 μM FtsZ to visualize FtsZ filaments. Grids (Ted Pella, Inc.) were glow discharged before use. Before preparing the grids, each sample was incubated for 10 minutes in the presence of 1 mM GTP at 30 ºC to allow for adequate assembly. Each sample was stained three times with 2% uranyl acetate for 20-seconds each. Each staining involved wicking the solution away and waiting 10 seconds in between stains. Samples were visualized using an JEOL JEM-1400 120kV Transmission Electron Microscope (TEM). Images were adjusted in contrast and brightness.

**GTPase Activity Assays**: GTP hydrolysis activity was measured using a continuous, regenerative coupled GTPase assay (8). In 96-well plates, 5 μM FtsZ in Buffer MES was used, supplemented with 1 mM phosphoenolpyruvate, 250 μM NADH, and 40 units $ml^{-1}$ of pyruvate kinase/lactic dehydrogenase from rabbit muscle (Millipore Sigma) at 30 ºC. A linear decline of NADH absorbance at 340 nm was monitored over 30 minutes using a SpectraMax i3x Multi-Mode Microplate Reader (Molecular Devices). Rate of GTP hydrolysis was calculated from the raw data of $A_{340}$ min$^{-1}$ and normalized to the hydrolysis rate of WT FtsZ.

**UV-CD measurements.** CD measurements were carried out using a Jasco 810 spectropolarimeter scanning from 260 nm to 190 nm, with a data pitch of 1 nm and a bandwidth of 1 nm. Four to six accumulations were averaged for each spectrum with a scanning speed of 50 nm/min and a 2-s response time (9). Additional details are described in the *SI Appendix*. The [θ] of CTT, and accordingly, the [θ] of ΔCTT+CTT were rescaled to be on the similar scale as that of WT and ΔCTT due to differences in concentration determination of CTT by absorbance and ΔCTT and WT by Pierce Protein assay. This is indicated by an asterisk on the y-axis in **Figure 1D**.

A quartz cuvette with a path length of 1 mm was used for all measurements. Buffer MES (50 mM MES, 50 mM KCl, 2.5 mM $MgCl_2$, 1 mM EGTA, pH 6.5) filtered with a 0.2 μm PES filter was used in all measurements at 30 ºC. CD spectra were converted from machine units (mdeg) to mean residue ellipticity ([θ], deg cm$^2$ dmol$^{-1}$ residue$^{-1}$) as described in the recent work of Seim et al., (10). Peptide concentrations were determined by absorbance using an Implen NP80 nanophotometer. Molar concentration values were calculated using an extinction coefficient of 195 M$^{-1}$ cm$^{-1}$ at 258 nm.

**Deconvolution of CD data.** CD data sets were converted to units of mean residue ellipticity as previously described (10) and submitted to the Dichroweb server (11) (http://dichroweb.cryst.bbk.ac.uk) for deconvolution using the CDSSTR algorithm (12) using basis sets 4, 7, and SP175t, which are optimized for the wavelength range of 190-240 nm (13, 14). We present the α-helical, β-strand, and β-turn categories each as a sum of two sub-categories that are given in the output of the analysis. The plotted secondary structure content estimates represent the mean and standard deviation of the values results obtained from the three basis sets.

**Fluorescence Correlation Spectroscopy (FCS):** Synthesized CTT peptides labeled with tetramethylrhodamine (TMR) labels were used (WatsonBio). Peptides were thawed in containers with desiccant while the surfaces of the 8-well plates (0.17 ± 0.005 mm thickness) were passivated by adding 150 μL of BSA (2 mg/ml) to the wells and letting it sit for 10-15 minutes. After 15 minutes, the surfaces were washed 3-5x with 1 mL of DI water. Three stocks were generated by dilution in Buffer MES, assumed to have the same viscosity as water: (1) 50 nM peptide, (2) 5 nM peptide, (3) 5 nM free dye. The free dye was used to determine the hydrodynamic radius of the beam and used to calculate the hydrodynamic radius ($R_h$) of the peptides per the Stokes-Einstein Equation (15). The diffusion coefficient of TMR with the instrument setup used here is $4.30 \times 10^{-10}$ m$^2$s$^{-1}$. All data were collected on a Confocor II LSM system (Carl Zeiss-Evotec, Jena, Germany) with a 40x water-immersion objective. Samples were excited at 500 nm, and emission was collected at 620 nm. Data for fluorescence intensity autocorrelation functions were analyzed with Zeiss Confocor II FCS software.

**All-Atom Simulations of CTT Sequence Variants:** All-atom Monte Carlo simulations were performed using the ABSINTH implicit solvent model and forcefield paradigm as made available in the CAMPARI simulation package (http://campari.sourceforge.net) (16, 17). Simulations utilized the abs_3.2_opls.prm parameter set in conjunction with optimized parameters for neutralizing and excess $Na^+$ and $Cl^-$ ions (18). Each of the simulations were performed using spherical droplets with a diameter of 120 Å for isolated peptides and 200 Å for full-length proteins with explicit ions to mimic a concentration of 10 mM NaCl. For isolated peptides, temperature replica-exchange Monte Carlo (T-REMC) (19) was utilized to improve conformational sampling. The temperature schedule ranged from 280 K to 400 K. Ensembles corresponding to a temperature of 340 K were used in the analysis reported in this work. Three independent sets of T-REMC simulations were performed for each Kappa variant. In all, the ensembles for each Kappa variant were extracted from simulations, where each T-REMC simulation deploys $5.5 \times 10^7$ Monte Carlo steps. In each simulation, the first $5 \times 10^6$ steps were discarded as equilibration. For full length proteins, we used SWISS-MODEL (20, 21) to create a homology model of the *Bs* core from PDB ID: IW5B. The core was initially equilibrated to add any missing atoms from the PDB file and make sure bond lengths were consistent with the ABSINTH model. Following the initial equilibration of the core, a single simulation step was performed at 20 K to build in each CTT. Then the end PDB structure from this step was used to generate five distinct full length starting structures. Specifically, the core was frozen such that only side chain moves were allowed and then 20,000 simulation steps at 500 K were performed. The end PDBs from the five simulations were then used as the input structures for the five independent full simulations ran at 340 K. For each full simulation a total of $6.15 \times 10^7$ steps were performed with $10^7$ discarded as equilibration. Simulation results were analyzed using the MDTraj (22) and CAMPARITraj routines that are available at http://pappulab.wustl.edu/CTraj.html.

**Shannon Entropy Calculations:** Shannon entropy values were quantified as previously described (23). Here, only two- parameter distributions were used with $R_g$ and δ*. $R_g$ was divided by $n^{0.5}$ where n is the number of residues in each CTT sequence. This accounts for the difference in number of 4 residues between CTT variants and the WT-CTT. For each sequence-specific distribution, we tiled the shape (δ*) - and size ($R_g$) -axes into four evenly sized regions, giving rise to a total of 16 bins. The boundaries for each of the bins were computed using the maximum and minimum observed values across all variants.

### Table S1

| Variant | Sequence | $\kappa_{+-}$ |
|---|---|---|
| K14 | GS HQPKPEQKSEANQSREQVTRELHSRNVPEIQKKDKVPQQPSTNTSTPQRI PG-***CTP*** | 0.136 |
| K18 | GS VQNKQIEKQKEPERRQRTHVESSPHQPSSQPVDRNPLPKTNQTQEKASIT PG-***CTP*** | 0.182 |
| K20 | GS IKQERKVTKPQEPSLNQSIRTHNQSVPEKEPDERRPQQQNTVSKHTSQPA PG-***CTP*** | 0.197 |
| WT | IEQEKDVTKPQRPSLNQSIKTHNQSVPKREPKREEPQQQNTVSRHTSQPA PG-***CTP*** | 0.197 |
| K26 | GS IEQKEKVTKPQRPSLNQSIETHNQSVPRRKPREKDPQQQNTVSEHTSQPA PG-***CTP*** | 0.258 |
| K33 | GS KQSTNTPQEKQPSIKVRRQQEVKPSLQKQENTVHIRRHSNAPTSDPPEEQ PG-***CTP*** | 0.331 |
| K34 | GS HQKPRQKVNKRQSEIRVPQSELSRSPTQENEEQSQPPAKTKNVQDTIHTP PG-***CTP*** | 0.332 |
| K41 | GS HPPEQQISKTRKRHVTQSQQEDEPPAKQKSLRRQNQEINVVNSSKPETTP PG-***CTP*** | 0.408 |
| K47 | GS PQIHQPKPQKKRSSNPTDQHKSAVTTPRKRVLIRQQPQTESEVEEESNQN PG-***CTP*** | 0.466 |
| K53 | GS IEQEEDVTEPQEPSLNQSIRTHNQSVPKKKPRKKRPQQQNTVSRHTSQPA PG-***CTP*** | 0.528 |
| K59 | GS PIVLEEDEETSEVSTNQKTQQVKPPPINSSQPQQHHPQRAQRKRRKKTSN PG-***CTP*** | 0.591 |
| K72 | GS NSQTQIRKRRKKKKRSSHQIVNLPNAPPDPEEEESEPHQSQTQPTVTQVQ PG-***CTP*** | 0.717 |

**Table S1: Sequences of the Kappa variants with the conserved CT17 motif shown as *CTP*.** We designed ten CTT sequences with $\kappa_{+-}$ values spanning from 0.14 to 0.72. Column 1 shows the name assigned to each *B. subtilis* FtsZ variant; columns 2 and 3 show the designed sequence of the CTL and the $\kappa_{+-}$ value for the variant, respectively. K33 is identical to previously reported Scr construct (24). Sequences of Kappa variants are 4 residues longer because of the cloning artifacts (see Methods).

### Table S2: List of plasmids and strains used in the study

| Strain or Plasmid | Genotype/features | Source |
|---|---|---|
| | ***B. subtilis*** | |
| JH642 | trpC2 pheA1 | Perego et al 1988 |
| PAL 2084 | JHC642 thrC::$P_{xyl}$-ftsZ | Weart Levin 2003 |
| PAL 3171 | PAL 2084 amyE::$P_{spac}$-ftsZ | Buske Levin 2012 |
| PAL 3491 | PAL 2084 *amyE*::$P_{spac}$-*ftsZCTLV1* | This study |
| PAL 3495 | PAL 2084 *amyE*::$P_{spac}$-*ftsZCTLV2* | This study |
| PAL 3499 | PAL 2084 *amyE*::$P_{spac}$-*ftsZCTLV3* | This study |
| PAL 3503 | PAL 2084 *amyE*::$P_{spac}$-*ftsZCTLV4* | This study |
| PAL 3507 | PAL 2084 *amyE*::$P_{spac}$-*ftsZCTLV5* | This study |
| PAL 3511 | PAL 2084 *amyE*::$P_{spac}$-*ftsZCTLV6* | This study |
| VK 1 | PAL 2084 *amyE*::$P_{spac}$-*ftsZK19* | This study |
| VK 2 | PAL 2084 *amyE*::$P_{spac}$-*ftsZ*K53 | This study |
| VK 3 | PAL 2084 *amyE*::$P_{spac}$-*ftsZK59* | This study |
| | ***E. coli*** | |
| AG1111 | *DZR200- MC1061 F'lacIQ lacZM15 Tn10 (tet)* | Ireton et al 1993 |
| PAL 930 | AG1111 + pBS58 | Weart Levin 2003 |
| **Plasmids** | | |
| pPJ19 | pET21b(+)-ftsZ946-51 Bam HI 1090-95 XmaI stop | Buske Levin 2013 |
| pPJ53 | pET21(+)-*ftsZCTLV1* stop | This study |
| pPJ54 | pdR67-*ftsZCTLV1* stop | This study |
| pPJ55 | pET21(+)-*ftsZCTLV2* stop | This study |
| pPJ56 | pdR67-*ftsZCTLV2* stop | This study |
| pPJ57 | pET21(+)-*ftsZCTLV3* stop | This study |
| pPJ58 | pdR67-*ftsZCTLV3* stop | This study |
| pPJ59 | pET21(+)-*ftsZCTLV4* stop | This study |
| pPJ60 | pdR67-*ftsZCTLV4* stop | This study |
| pPJ61 | pET21(+)-*ftsZCTLV5* stop | This study |
| pPJ62 | pdR67-*ftsZCTLV5* stop | This study |
| pPJ63 | pET21(+)-*ftsZCTLV6* stop | This study |
| pPJ64 | pdR67-*ftsZCTLV6* stop | This study |
| pVK1 | pET21(+)-*ftsZK19* stop | This study |
| pVK2 | pdR67-*ftsZK19* stop | This study |
| pVK3 | pET21(+)-*ftsZK53* stop | This study |
| pVK4 | pdR67-*ftsZK53* stop | This study |
| pVK5 | pET21(+)-*ftsZK59* stop | This study |
| pVK6 | pdR67-*ftsZK59* stop | This study |

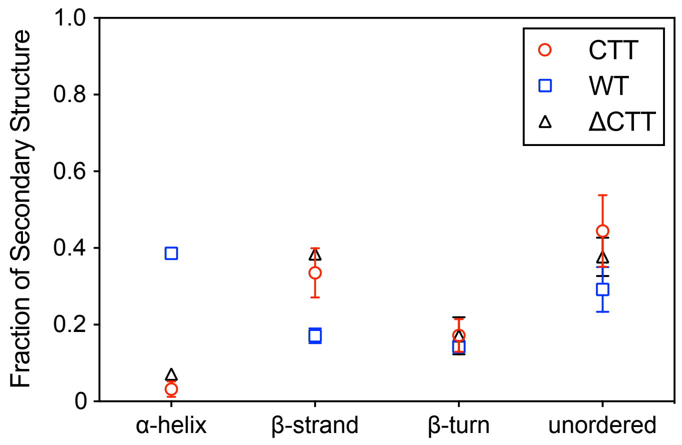


**Figure S1. Secondary structure contents of each construct extracted from CD measurements.** Secondary structures were determined by deconvolution using the CDSSTR algorithm and basis sets 4, 7, and SP175t, as implemented by the Dichroweb server. Open circles and error bars represent the mean and standard deviation, respectively, of the analysis resulting from these three basis sets.

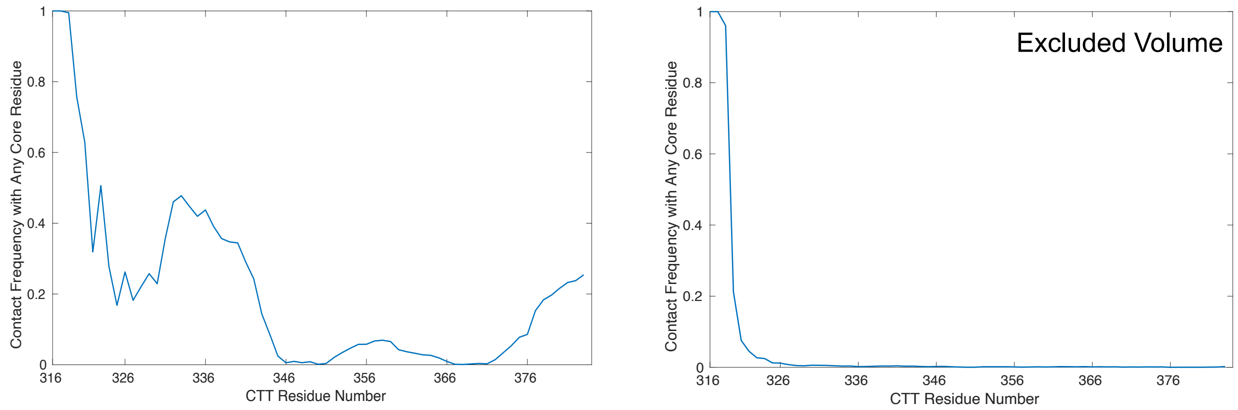


**Fig. S2. Frequency of CTT residues making contacts with any residue on the core domain.** (A) This panel shows the results obtained using the default parameters of simulations. This result is mapped onto the *Bs*-FtsZ structure and shown in **Figures 2A and 2C** in the main text. (B) In a stark contrast, when the CTT in the FtsZ monomer was modeled as a self-avoiding polymer, retaining the intrinsic flexibility of the CTT sequence and turning off all specific interactions other than those to steric exclusion, the CTT does not interact with the core domain except for the few N-terminal residues that are immediately proximal to where the CTT is tethered to the core. This highlights the specificity of the contacts we map using the atomistic simulations.

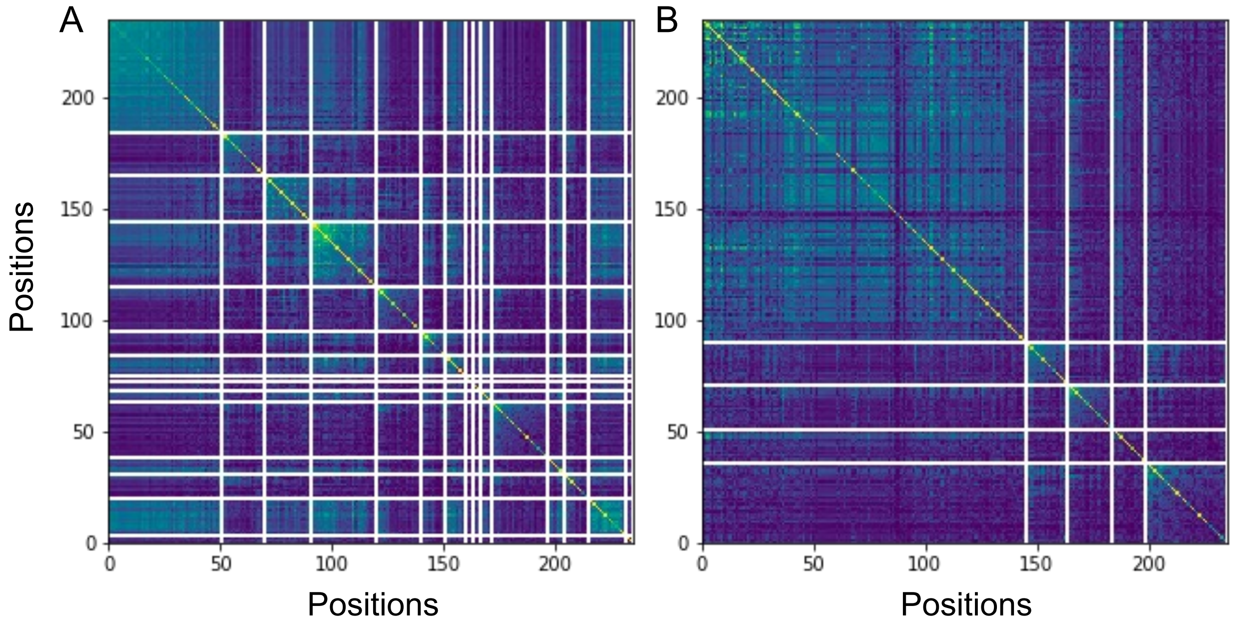


**Figure S3. Covariance matrices of FtsZ orthologs obtained from statistical coupling analysis.** (A) Fifteen independent components (ICs) were determined. (B) ICs with strong inter-group correlations were combined to yield a total of five sectors.

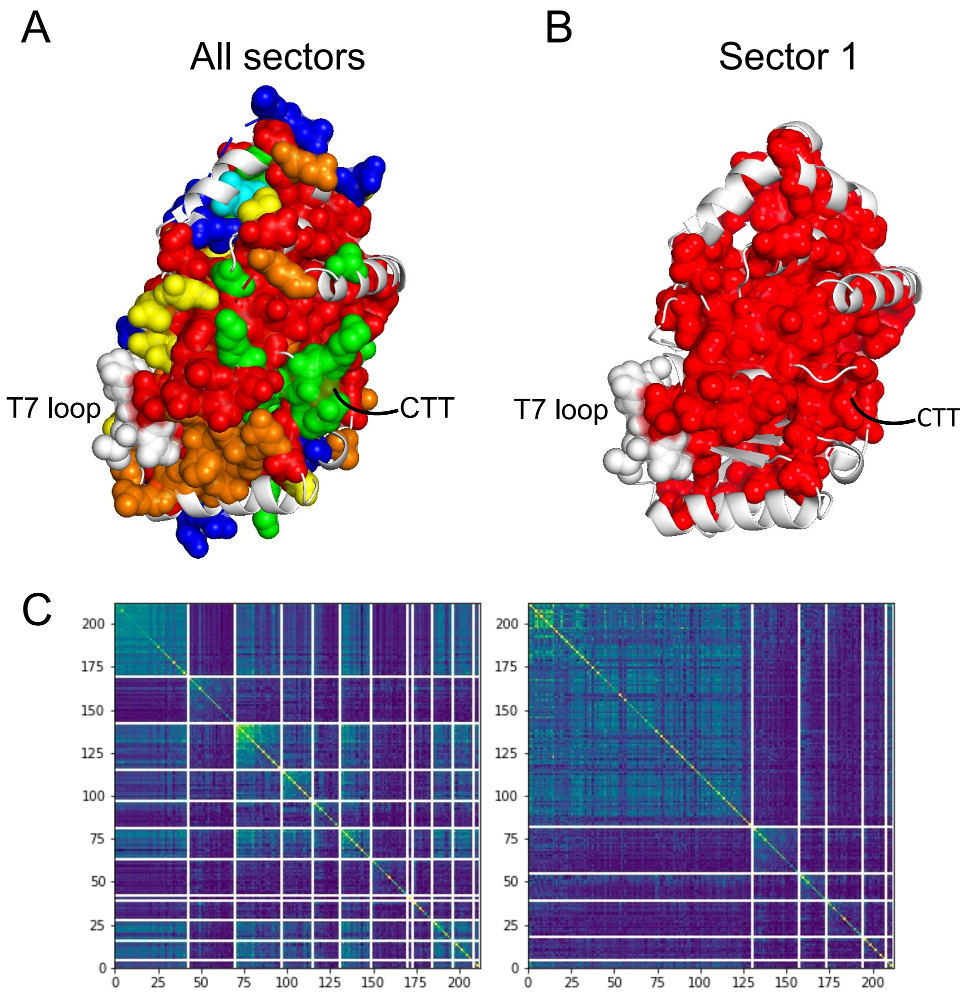


**Figure S4: Identified sectors can be mapped onto the core domain of Bs-FtsZ**. (A) Six sectors were identified by SCA, and each sector is color coded (left) in red, orange, yellow, green, blue, and cyan (25). The residues that are not part of any sectors are shown in white. The T7 loop is shown in space filling model and is not part of any sectors (white) when only the core domain is used for multiple sequence alignments. (B) Only Sector 1 is shown for clarity. (C) Covariance matrices of core domains from SCA show 12 ICs (left) that yield a total of six sectors (on the right).

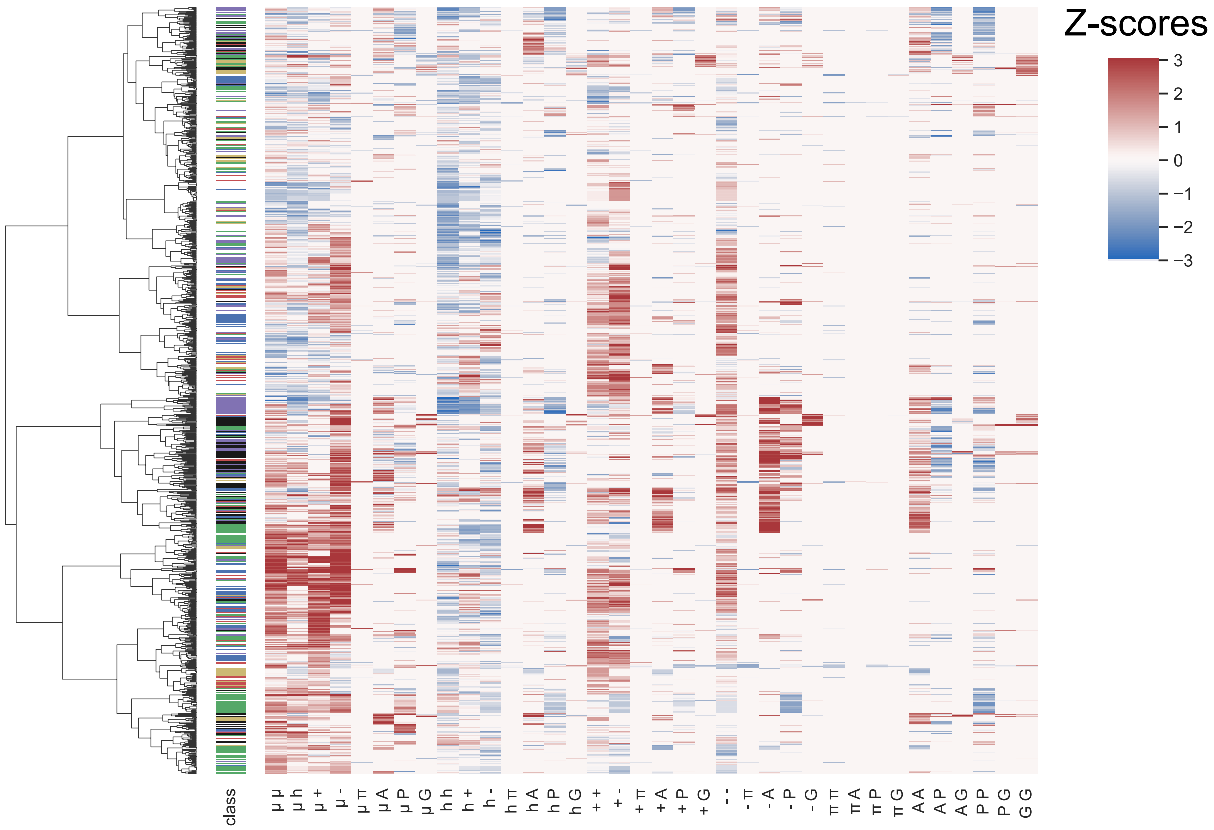


**Figure S5: Comparison of linear sequence patterns of the CTTs from 1208 FtsZ orthologs.** Each row below the "class" row is a heatmap of z-scores for a distinct binary pattern. Each column represents a sequence. Conserved non-random features refer to binary patterns that have non-zero z-scores across most orthologs. Corresponding class assignments for each sequence are shown in the "class" row and color coded as noted below. The dendrogram shows the similarity of z-score matrices across orthologs. The dendrogram was generated using the Frobenius norm of the z-score matrices, where the norms were used as Euclidean distances, and Ward's clustering was used. The lack of clustering by classes indicates that the sequence patterning features are not phylogenetically correlated. The row labels describe the pair of residues or residue types for which the z-scores were calculated: μ = polar, h = hydrophobic, + = positively charged, - = negatively charged, π = aromatic, A = alanine, P = proline, G = glycine. The class column indicates phylogenetic classes as follows: *clostridia* = red, *gammaproteobacteria* = green, *bacilli* = blue, *actinobacteria* = black, *alphaproteobacteria* = magenta, *betaproteobacteria* = yellow.

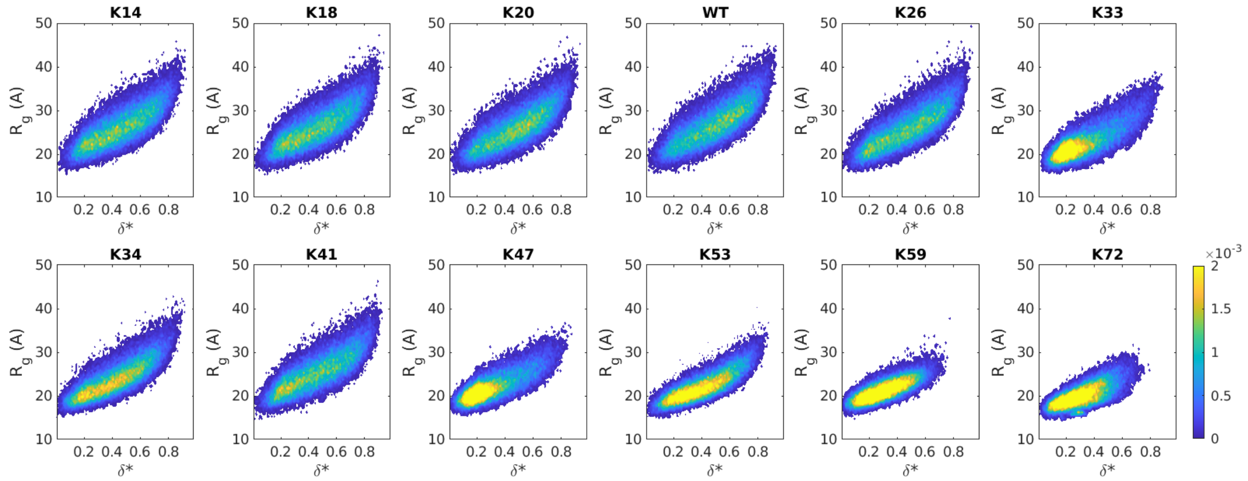


**Figure S6: Conformational distributions of isolated CTT peptides of Kappa variants.** The figure shows probability density distributions of asphericity (δ*) and $R_g$ obtained from simulations. The color bars show the probability densities. The bin widths along the abscissa and ordinate are 0.01 and 0.5 Å, respectively. Data are shown for simulations performed at a temperature of 340 K. Low values of δ*correspond to globules, intermediate values of δ* correspond to prolate ellipsoids, and values of δ* tending to unit are rod-like conformations. For CTT peptides with $\kappa_{+-} \leq 0.26$, the distributions are relatively flat across the asphericity axis, spanning the range of δ* values from 0.1 – 0.8 and $R_g$ axis ranging from ~ 17 Å – 42 Å. The probability density associated with compact, globular conformations increases as $\kappa_{+-}$ increases above 0.33.

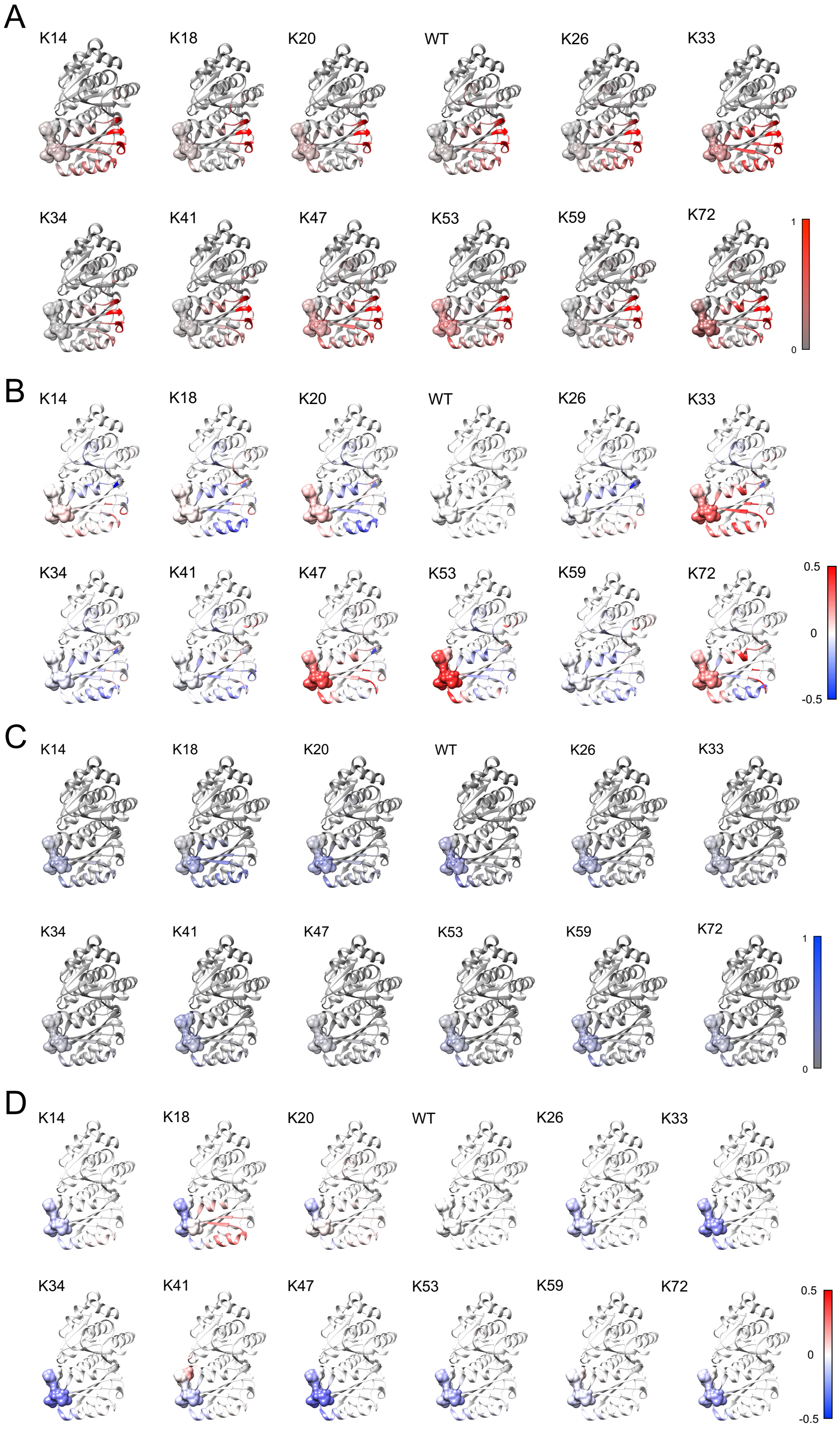


**Figure S7: Intra-molecular interactions between the core and CTL or CT17 in Kappa variants.** The colors on the structures of *Bs*-FtsZ core domain indicate the fractional frequency of (A) CTL or (C) CT17 coming within 10 Å of the residues in the core domain as indicated in the color bar on the right. Differences in fractional frequency of (B) CTL-core or (D) CT17-core interactions between each Kappa variant and WT are mapped onto the structure of the core domain.

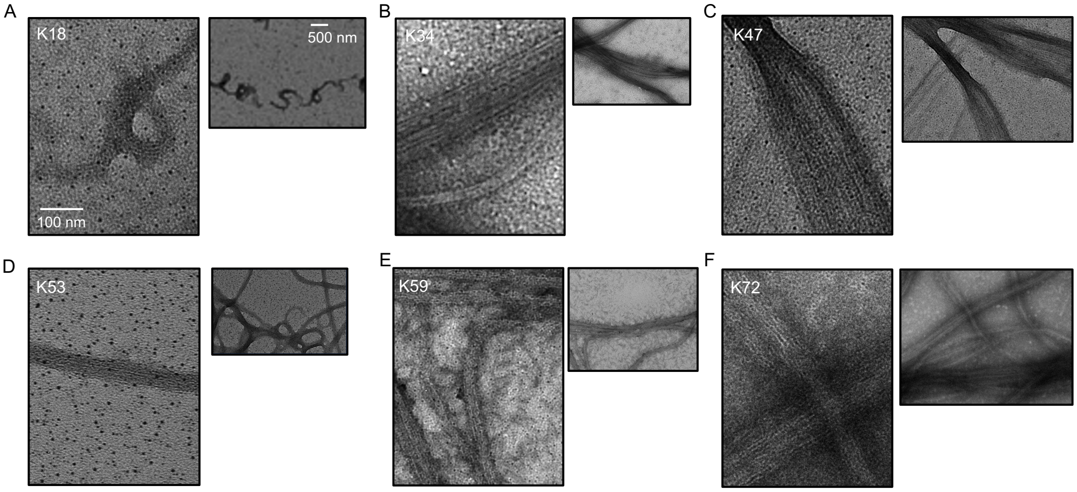


**Figure S8: TEM images of select Kappa variants**. The Kappa variants presented here were chosen to highlight the non-native, large assemblies formed by what appear to be tail-mediated interactions. Structures. The images are shown using scale bars of 100 nm and 500 nm. These scale bars help highlight show the large size of the assemblies and the lateral associations of filaments.

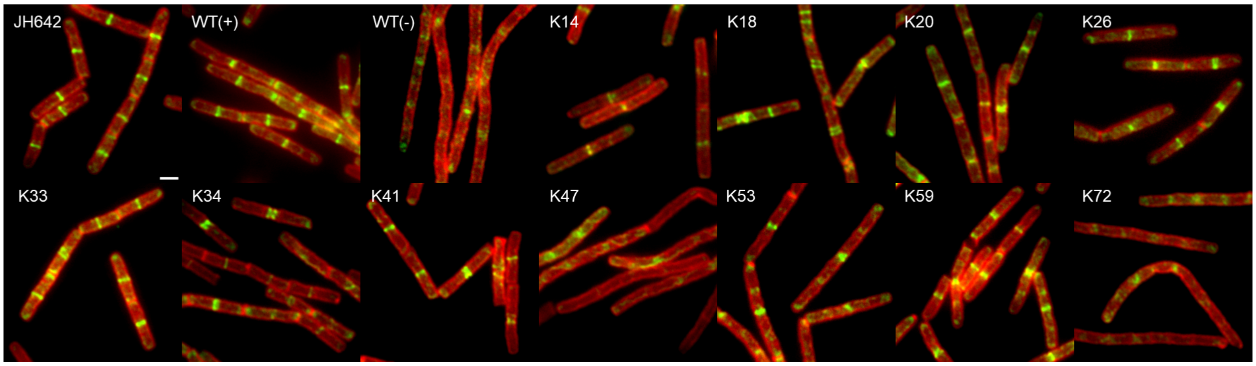


**Figure S9: IFM images of representative *B. subtilis* cells expressing Kappa variants.** The imaged cells were artificially colored in green for FtsZ and red for the cell wall. The cell septa can be identified by high intensity of red lines between two or more *B. subtilis* cells at their poles. Scale bar = 2 µm.

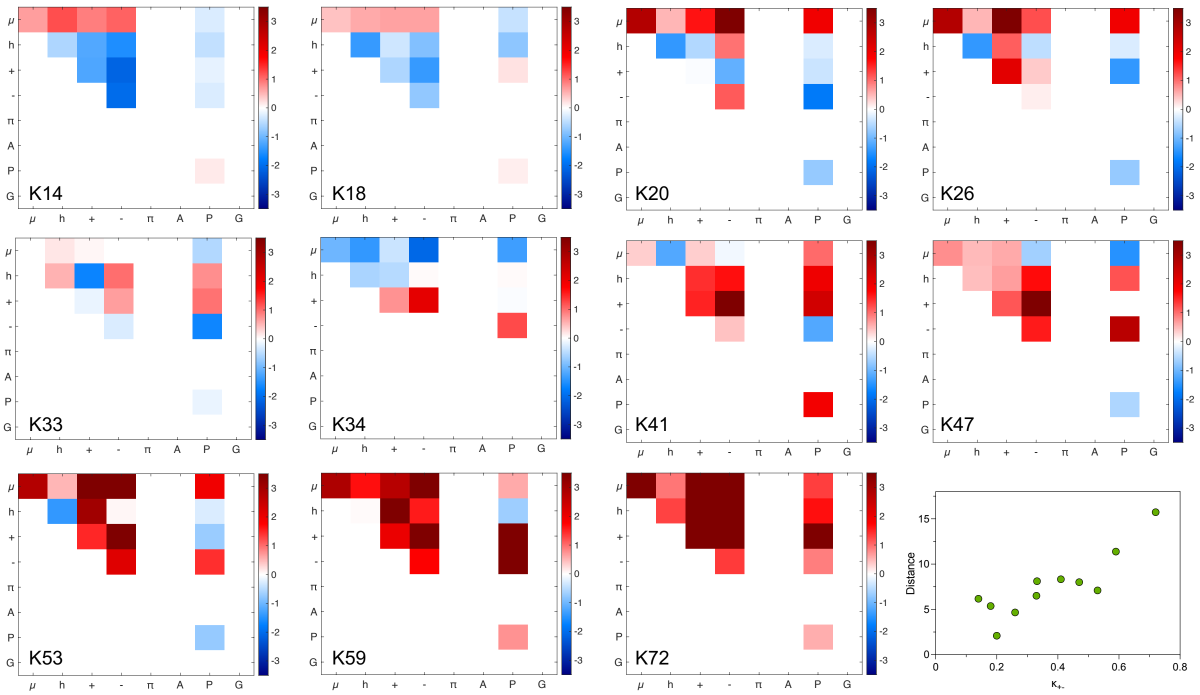


**Figure S10: Z-score matrices of CTT sequences for Kappa variants.** The data show results from NARDINI analysis. The Euclidean distance between the z-score matrices of the WT CTT and CTTs of Kappa variants show a non-monotonic variation with $\kappa_{+-}$.
